# Proteomic profiling of the neuroblastoma secretome identifies extracellular vesicles as drivers of T cell suppression

**DOI:** 10.64898/2025.12.02.690421

**Authors:** Josephine G. M. Strijker, Ronja E. van Berkum, Elisavet Kalaitsidou, Arjan Boltjes, Naima Hiddink Verberne, Mirjam A. Damen, Femke van den Ham, Liselotte E. Baaij, John Anderson, Jan J. Molenaar, Wei Wu, Judith Wienke

## Abstract

Immunosuppressive tumor microenvironments in solid tumors hamper the efficacy of immunotherapies, including CAR T-cell therapy. To identify immunosuppressive secreted factors in high-risk neuroblastoma, a childhood solid tumor with a 5-year overall survival of below 60%, we mapped secreted factors of 16 genetically diverse patient-derived neuroblastoma tumoroids by LC-MS. The secretomes suppressed T cell activation, proliferation and cytotoxicity to varying extents, suggesting that this panel constituted a valuable discovery foundation to identify clinically relevant immunosuppressive factors. 29 proteins were significantly enriched in highly suppressive secretomes, of which Neuropeptide Y (NPY) had the highest correlation with functionally-determined immunosuppression. No direct or indirect effect of NPY on (CAR) T cell activation could be validated. Instead, NPY proved to be a biomarker for extracellular vesicle (EV) secretion. Finally, we demonstrated that neuroblastoma-derived EVs potently suppress T cell activation. Altogether, this study provides an atlas of the neuroblastoma secretome and reveals immunosuppressive effects of EVs on T cells. These insights contribute to the understanding of the immunosuppressive neuroblastoma TME.

## Introduction

Immunotherapy has revolutionized cancer therapy, and has provided encouraging results in cancers with previously poor prognoses^1,2^. Adoptive cell therapies, e.g. chimeric antigen receptor (CAR) T-cells, re-program the patient’s own immune cells to attack cancer cells. CAR-T cells have been particularly successful in hematological malignancies^3,4^. For solid tumors, however, the efficacy of CAR T-cells still remains limited. Multiple factors have been identified to contribute to the failure of CAR-T cells in solid tumors, including the immunosuppressive tumor microenvironment (TME)^5^. Immunosuppressive factors in the TME include immune checkpoint expression and the secretion of immunomodulatory factors^5^. Dissecting how solid tumors modulate the TME into an immunosuppressive environment is pivotal to optimize the efficacy of CAR T-cells and other immunotherapies.

Neuroblastoma is an extracranial pediatric solid tumor, arising from sympathetic nervous cells, which often metastasizes to the liver and bone marrow^6^. Especially the high-risk group, which includes patients with a higher tumor burden and specific genetic alterations such as MYCN amplification, have a poor prognosis, despite an intense multi-modal treatment regimen^7,8^. Previous clinical CAR T-cell trials in high-risk neuroblastoma showed limited clinical efficacy, with suboptimal CAR T-cell expansion and persistence^9–11^. A more recent clinical trial with a third-generation CAR T-cell targeting GD2 reported promising results with an overall response rate of 63% and improved CAR-T cell persistence^12^. However, clinical responses remained limited to patients with a low disease burden. The inadequate efficacy of CAR T-cells, especially in patients with a high tumor burden, suggests that immunosuppressive factors in the TME may hamper CAR-T cell responses.

The cancer secretome – all soluble factors and extracellular vesicles (EVs) secreted by cancer cells – is still a relatively under-studied element of the TME, although it is thought to play a significant role in cell-to-cell communication within the TME^13^. EVs are secreted membrane-bound vesicles released by cells into the extracellular space, where they interact with other cells and can play a role in immune regulation^14,15^. The cancer secretome can drive tumor proliferation, chemotherapy resistance, metastasis, and immunosuppression both locally and systemically^16–20^. For instance, a well-known tumor-derived soluble factor is Tumor Growth Factor-β (TGF-β), which can suppress cytotoxic and helper lymphocytes while activating suppressive regulatory T cells^21,22^. Despite promising preclinical evidence, clinical success to enhance immunotherapy by blocking TGF-β remains limited^23^. For neuroblastoma specifically, we recently demonstrated that blocking immunosuppressive tumor-secreted Macrophage Migration Inhibitory Factor (MIF) enhances CAR T-cell efficacy *in vitro* and *in vivo*^24^. Next to soluble factors, tumor cells secrete EVs, influencing recipient cells by transferring their cargo and modulating cellular functions^25^. EVs can, for example, express checkpoints, such as PD-L1 or CD80, or carry tumor metabolites such as lactate to directly suppress T cell activation and longevity^26–28^. A comprehensive map of the neuroblastoma secretome is still lacking but could reveal new and improved targets to enhance immunotherapy efficacy.

Here, we present an unprecedented proteomic atlas of neuroblastoma secretomes which identifies a total of 1686 tumor-derived proteins. We identify 29 candidate proteins associated with suppression of T cell activation. In addition, pathway analysis along with *in vitro* exposure assays indicate that extracellular vesicles (EVs) drive this immunosuppression. These data contribute to a deeper understanding of the immunosuppressive TME in neuroblastoma and aid the future development of tailored immunotherapies.

## Material and Methods

### Tumoroid culture

Patient derived tumoroid cultures were hestablished from fresh biopsy or resection neuroblastoma tumor material, obtained as part of the ‘iTHER’-study or the biobank initiative of the Princess Máxima Center for Pediatric Oncology, Utrecht, Netherlands, or were established previously^29–31^. Ethics approval was granted for all samples: patients and/or their legal representatives signed informed consent, and the study was conducted in accordance with the Declaration of Helsinki. In short: tumors were mechanically and enzymatically digested, washed and placed in optimal culture conditions (37°C, 5% CO_2_) in optimized medium (DMEM, low glucose, GlutaMAX supplemented with 20% F-12 Ham’s Nutrient Mix, 100U/mL penicillin, 100µg/mL streptomycin, B27 (50x), N2 (100x), hIGF (300ng/mL), hFGF (40ng/mL), hEGF (20ng/mL), PDGFaa (10ng/mL), PDGFbb (10ng/mL)). A panel of successful long-term neuroblastoma tumoroid cultures was chosen for this project and cultured to be expanded to full T175 flasks. Cells were passaged and medium was refreshed twice a week.

### Secretome harvest, sample preparation and LC-MS

Two T175 flasks of tumoroids were washed and incubated with 25 mL empty DMEM GlutaMAX for 24h. The 50 mL conditioned medium was concentrated to 1 mL using 3kDa molecular weight cut-off filters (Millipore). Protein concentration was measured via DC protein assay (Bio-Rad). Aliquots for functional assays were stored at -70°C. Aliquots for LC-MS were denatured in 4 M urea (pH 8.0) and stored at -70°C. Proteins were reduced with DTT (20°C, 1 h), alkylated with IAA (20°C, 30 min, dark), diluted to <2 M urea, and digested with Lys-C (4 h, 37°C) and trypsin (overnight, 37°C). Peptides were desalted (Sep-Pak C18), vacuum-dried, and stored at -80°C. Peptides were reconstituted in 2% formic acid and were analyzed in triplicate on an Orbitrap Exploris 480 mass spectrometer connected to an UltiMate 3000 UHPLC system. Peptides were trapped on a µ-precolumn (C18 PepMap100, 5 µm, 100 Å, 5 mm × 300µm; Thermo Scientific) and separated on an analytical column (120 EC-C18, 2.7 µm, 50 cm × 75 µm; Agilent Poroshell) using a gradient of 0-10% solvent B (10 min), 10-40% (138 min), and 100% (2 min), followed by re-equilibration.

### Raw secretome data analysis

Raw spectral files were searched with MaxQuant (version 2.1.4.0) against the human UniProt database (downloaded in September 2022) using the integrated Andromeda search engine. Cysteine carbamidomethylation was included as a fixed modification, while protein N-terminal acetylation and methionine oxidation were allowed as variable modifications. Label-free quantification (LFQ) algorithm and the match-between-runs feature were enabled for protein identification. Trypsin/P was set as the digestion enzyme (cleaves after lysine, even if a proline follows), and up to two missed cleavages were allowed. A false discovery rate (FDR) was restricted to 1% for both protein and peptide identification. Bona fide secreted proteins were determined by referencing SignalP (version 5.0) for presence of classical signal peptides and Exocarta (version 2015) for non-classical but curated secretion.

### Suppression assays

Peripheral blood mononuclear cells (PBMCs) were isolated from healthy donor whole blood using Ficoll-Paque (Sigma-Aldrich) density centrifugation and stored in liquid nitrogen. PBMC vials were thawed and stained with anti-CD3-AF700 (Biolegend, #300324, 1:400) and fixable viability dye efluor506 (Invitrogen, #65-0866-14, 1:500) for FACS sorting of live CD3^+^ T cells on a Sony SH800S machine. Sorted T cells were labelled with 2μM CellTrace Violet (Invitrogen, #C34557) and placed in an anti-CD3 (1µg/mL, OKT3, Miltenyi) and anti-CD28 (5µg/mL, 15E8, Miltenyi) pre-coated plate with RPMI 1640 (Gibco) containing 10% human serum and either the concentrated secretomes with a final protein concentration of 10µg/mL, or recombinant NPY-w (1153, Tocris) or NPY-3-36 (N9407-250UG, Merck). After a 4 day incubation period, cells were harvested, washed and duplicates were pooled before staining (fixable viability dye efluor506; Invitrogen, cat. 65-0866-14, 1:500; CD25-BV711, BD, #563159, 1:50; CD3-AF700, Biolegend, #300324, 1:200; Granzyme B-RY586, BD, #568134, 1:800). Cells were fixated between the surface staining and the intracellular staining step with a FOXP3 fixation kit (Invitrogen). Flow cytometry data were acquired on a Cytoflex LX (Beckman Coulter). Flow cytometry data were analyzed using Flowjo (version 10.9.0, LLC).

### ELISA

Protein concentrations of NPY in concentrated secretome were measured using a NPY-ELISA kit (DY8517-05, Bio-Techne). Concentrated secretomes were diluted to 500µg/mL total protein concentration to measure relative NPY levels.

### NPY-R antagonist treated secretome

To determine the potential autocrine effect of NPY on tumoroid secretomes, tumoroid MaxNB400 was treated with recombinant NPY-w (1153, Tocris) or NPY-3-36 (N9407-250UG, Merck) for three days. Each day, fresh NPY was added to the culture. After the three day treatment period, empty DMEM GlutaMAX was added to the cells for 24hrs and conditioned medium was harvested, concentrated and stored for functional assays as described above. Tumoroid MaxNB293 was treated with α-NPY1R (BIBO3304, SML2094, Sigma Aldrich, 1µM), α-NPY2R (BIIE0246, 7377/1, Bio-Techne, 1µM), or α-NPY5R (CGP71683, 21-991-0, Tocris Bioscience, 1µM) after which secretome was harvested.

### B7-H3 CAR T-cell production

B7-H3 targeting CAR T-cells were produced as published previously^24,32^. In short: retrovirus was harvested from TE9-CAR construct producing Phoenix-Ampho lines, kindly provided by the Anderson lab (University College London)^32^. PBMCs were activated in a two-step manner with α-CD3 (130-093-387, Miltenyi, 0.5µg/mL) and α-CD28 (130-093-375, Miltenyi, 0.5µg/mL) and IL-2 (130-097-748, Miltenyi). T cells were transduced by spinoculation in a RetroNectin (T100A, Tekada, 8µL/mL) coated plate.

### IncuCyte killing assay

Tumoroid MaxNB293 was dissociated into single cells using Accutase (Sigma Aldrich), plated and left to grow back into spheres for 48 hours before CAR-T cells were added. CAR-T cells were pre-treated with α-NPY1R (BIBO3304, SML2094, Sigma Aldrich, 1µM) for 24 hours. CAR-T cells were washed and added to the tumoroid plates at a 1:10 Effector Target ratio, with fresh NPY1R antagonist added. Killing was measured with a Caspase-3/7-Green (4440, Sartorius, 5µM) based readout. Plates were imaged every 2 hours using the IncuCyte® Live-Cell analysis system (Sartorius). Data were normalized to the tumoroid only condition.

### Isolation of extracellular vesicles

25 ml of empty DMEM Glutamax was added to three T175 flasks. After 24 hours, cells were assessed and counted. The conditioned medium (75 ml) was pooled and centrifuged at 250g for 5 min at 4°C to remove dead cells, followed by two spins at ∼4000g for 10 min at 4°C to remove debris and apoptotic vesicles. The medium was filtered (0.45 µm, Millipore) and concentrated to 500 µl using a 100kDa ultrafiltration unit (Amicon). EVs were isolated via size-exclusion chromatography (SEC) using a qEV original column (Izon Science, Gen 2). Fifteen 500 µl fractions were collected, with RPMI 1640 (Gibco™) as the eluent. Fractions 5-9, containing EVs, were aliquoted and stored at −80°C.

### Western Blot analysis for EV fraction selection

To identify the three most EV-enriched fractions, western blotting was performed on fractions 5-9. Samples were reduced with 5mM DTT, heated at 95°C for 5 min, and 12 µl was loaded onto a 4-20% gradient gel (BioRad). Neuroblastoma SH-SY5Y whole cell lysate (WCL) was used as a positive control. Gels were run at 125 V for 90 min and transferred to PVDF membranes using the Biorad Trans-Blot Turbo system. Membranes were blocked for 1h at room temperature with ECL Prime reagent (Cytiva) and probed with antibodies for CD81, CD9, Flotillin-1, and Calnexin (CST, #52892, #13403, #18634 and #2679, 1:1000 each). Chemiluminescence was detected using the Biorad ChemiDoc MP imager. Based on these results, the three most EV-enriched fractions were selected for further experiments.

### Nanoparticle tracking analysis (NTA)

NTA was performed to determine EV concentration and size distribution. The three EV-enriched fractions were pooled and diluted 1:50 in sterile PBS. Sterile RPMI diluted 1:50 in PBS served as a background control. EVs were analyzed using a NanoSight NS500 with an sCMOS camera, capturing three 30-second videos per camera level (12, 14, and 16). Analysis was conducted using NTA 3.4 software with a detection threshold of 5. The average particle concentration (particles/ml), after background subtraction, was calculated for further experiments.

### EV-exposure assay

To evaluate the effects of tumor-secreted EVs on T cell function, isolated EVs from NB tumoroids were added to activated healthy-donor T cells isolated in identical manner as described in ‘suppression assay’. Sorted T cells were labelled with 2μM CellTrace Violet (Invitrogen, #C34557) and placed in an anti-CD3 (1µg/mL, OKT3, Miltenyi) and anti-CD28 (5µg/mL, 15E8, Miltenyi) pre-coated plate with RPMI 1640 (Gibco) containing 10% human serum, in addition to 120µl EVs per well, isolated from tumoroid lines MaxNB067, AMC691B and MaxNB293, in a 96 well plate. After a 4 day incubation period, cells were harvested, washed and duplicates were pooled before staining (fixable viability dye efluor506; Invitrogen, cat. 65-0866-14, 1:300; CD25- APC/Fire810, Biolegend, #356149, 1:25; CD3-AF700, Biolegend, #300324, 1:400; Granzyme B- PE/dazzle 594, Biolegend, #372215, 1:100). Cells were fixated between the surface staining and the intracellular staining step with a FOXP3 fixation kit (Invitrogen). Flow cytometry data were acquired on a Cytoflex S (Beckman Coulter). Flow cytometry data were analyzed using Flowjo (version 10.9.0, LLC).

### Statistics

Statistical tests were performed using GraphPad Prism v10.0.2 software or R. Statistical tests are specified in the figure legends. A p-value of <0.05 was considered statistically significant, unless stated otherwise. To correct for multiple testing, the false discovery rate (FDR) was calculated and q-values are displayed. The differential expression analyses were performed with a Student’s t-test, assuming a 2-tailed distribution with equal variance. Correlation analyses were performed with a 2-tailed nonparametric Spearman correlation analysis.

## Results

### Patient-derived tumoroids to study the neuroblastoma secretome

To map the neuroblastoma secretome, i.e. the totality of constitutively secreted factors, we studied 16 established patient-derived tumoroids grown from primary or metastatic neuroblastoma tumors^33^. The tumoroids were cultured in empty medium for 24 hours, after which the conditioned medium was collected and concentrated for analysis by liquid-chromatography mass-spectrometry (LC/MS) and functional assays (Fig. 1A-B). We used a panel of 16 tumoroids from 14 patients with varying genetic aberrations, representing different time points of the disease course (Supplement Fig. 1A). From two patients (AMC700 and AMC691), tumoroids were available both from the primary tumor (annotated with T) and a bone marrow metastasis (annotated with B). 12 tumoroids had a MYCN amplification while 4 did not have a MYCN amplification (Fig. 1C, Supplementary Fig. 1A). Three non-malignant cultures were included as references: two fibroblast cultures grown from neuroblastoma tumors (MaxFIB293, matched with tumoroid MaxNB293, and MaxFIB272, matched with tumoroid MaxNB272) and HMEC-1, an immortalized microvascular endothelial cell line^34^.

**Figure 1:**
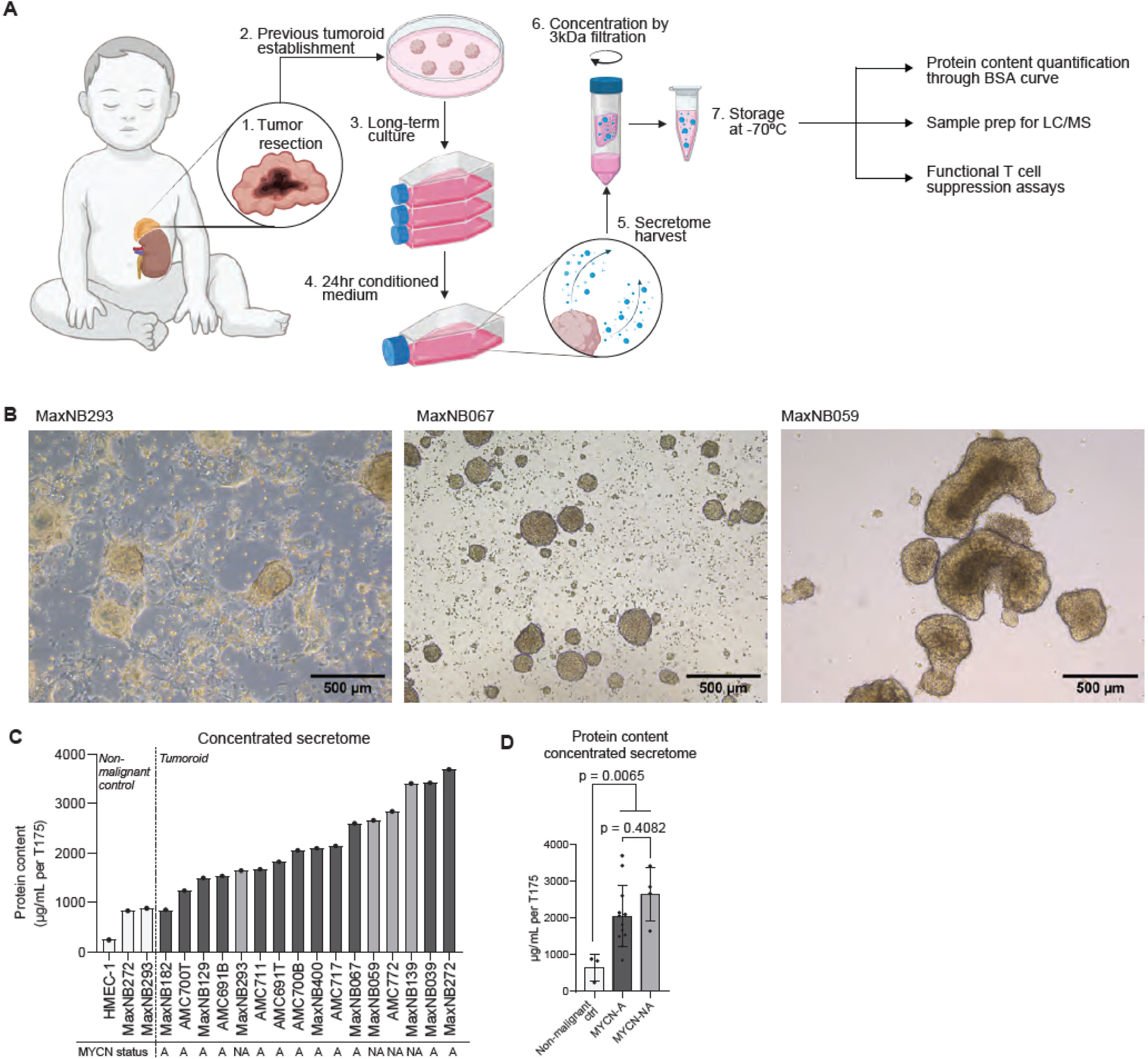
Patient derived tumoroids for secretome harvest **A:** Schematic representation of workflow for secretome harvest from patient derived tumoroids. Figure made with Biorender.com. **B:** Pictures from cultures of three representative tumoroid models. **C:** Protein concentration of non-malignant control and tumoroid secretomes as measured by Bradford protein assay. Total protein concentration was normalized to the number of full T175 flask harvested from. MYCN status of all tumoroid is indicated as A (Amplified) and dark grey or NA (non-amplified) and light grey. **D:** Comparison of protein concentrations of secretomes from non-malignant controls and tumoroids with MYCN-A or MYCN-NA. Statistics between non-malignant control and both tumoroid groups represents an unpaired t-test, while p-value for MYCN-A vs. MYCN-NA represents ordinary one-way ANOVA with Tukey’s multiple comparisons test.

Protein content of the concentrated secretomes was lowest in the non-malignant control samples (241.5 to 880 µg/mL per full T175), while the protein concentration of the tumoroid secretomes was significantly higher, ranging from 848.8 µg/mL to 3691 µg/mL for one full T175 flask, with the highest protein content in the secretome of tumoroid MaxNB272 (Fig 1C-D, Supplement Table 1). Protein concentration of the concentrated secretomes correlated with the total number of cells counted per flask (Supplementary Fig. 1B). No significant difference was observed between MYCN amplified and MYCN non-amplified tumoroids (Fig. 1D). To conclude, we harvested secretomes from these 16 diverse neuroblastoma tumoroids and 3 non-malignant references and proceeded to analyze the secreted factors in detail.

### A proteomic atlas of the neuroblastoma secretome

To dissect which factors are secreted by neuroblastoma tumoroids, we analyzed the secretome contents using liquid chromatography–mass spectrometry (LC-MS). The number of proteins found per line ranged from 340 to 1246. 61% - 88% were secretome annotated (SignalP iz + Exocarta), indicating a high confidence that they are bona fide secreted factors and not derived from the cytoplasm of dying or disintegrated cells (Fig. 2A, Supplementary Fig. 2A)^35–37^. Considering only proteins annotated as secretome-associated, 1686 unique proteins were detected in total from 19 tumor and non-malignant control samples (Fig. 2B).

**Figure 2:**
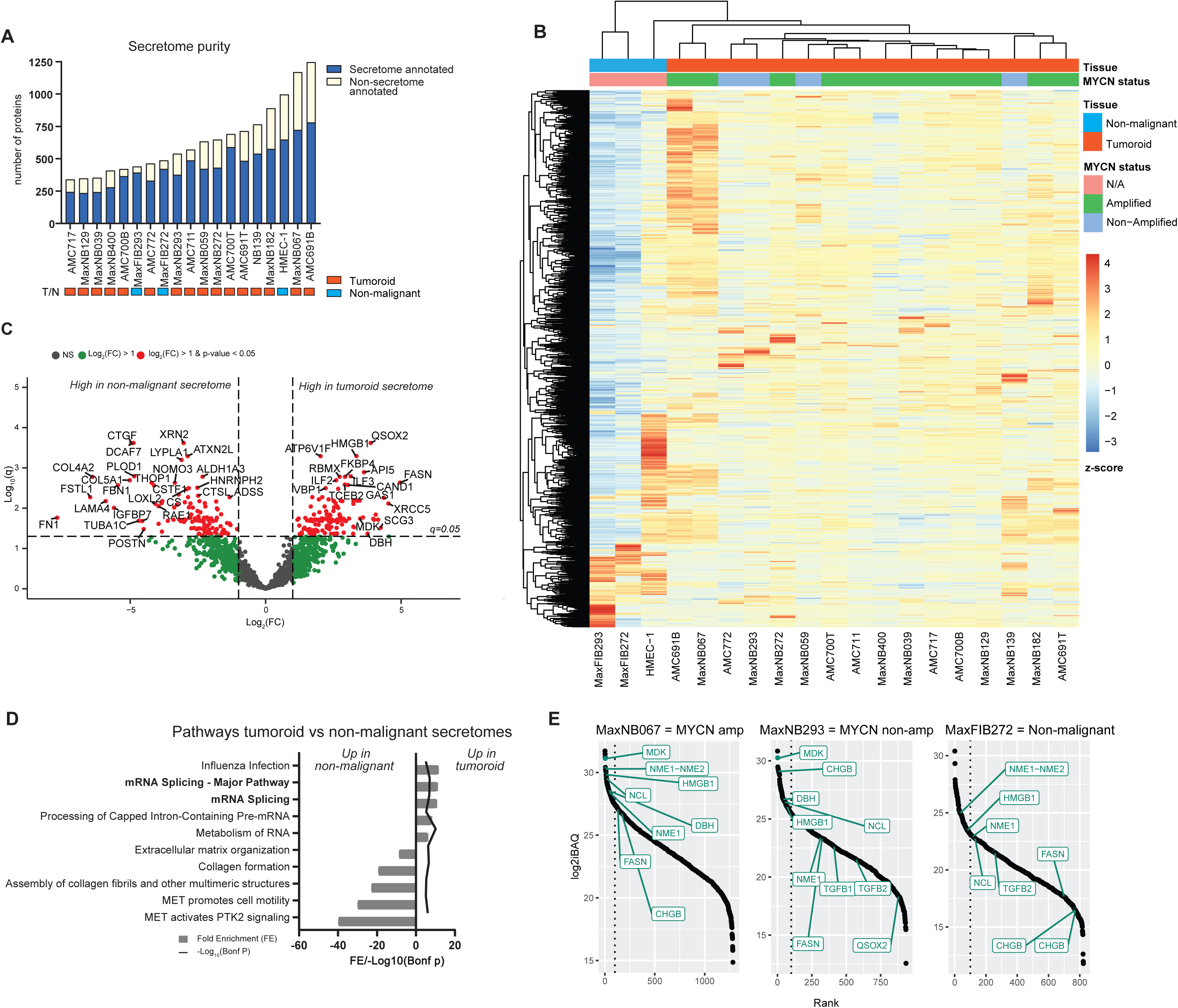
Map of secretome data. **A:** Bar plot indication the number of unique proteins identified per secretome. Secretome annotated proteins are annotated in blue. Non-secretome annotated are colored in yellow. **B:** Heatmap of z-scored imputed LFQ-intensities of all secretomes. Each column represents a tumoroid or non-malignant reference secretome. **C:** Volcano plot comparing all tumoroid secretomes to non-malignant control secretomes. **D:** Reactome pathways upregulated in normal secretomes versus tumoroid secretomes. **E:** Ranking of proteins in representative secretomes. iBAQ (intensity Based Absolute Quantification) indicates the protein’s non-normalized intensity divided by the number of measurable peptides and, hence, indicating the relative abundance of the protein in the sample. Vertical dotted line indicates rank 100. All factors mentioned in the main text are annotated.

From this rich dataset, we identified 129 secreted proteins which were significantly enriched in the tumoroid secretomes compared to non-malignant reference lines (Fold Change (FC) > 1, p-adjusted (p-adj) < 0.05) (Fig. 2C). Amongst these, several factors emerged of particular interest due to their potential relevance in tumor progression or TME regulation, including Fatty Acid Synthase (FASN), Quiescin Q6 sulfhydryl oxidase 2 (QSOX2) and Midkine (MDK). FASN (FC = +4.96, p-adj = 0.0023), which is reported to play a role in the regulation of tumor-metabolism and energy consumption, had the highest fold change^38^. QSOX2 (FC = +3.88, p-adj = 0.0002) was identified to regulate apoptosis in neuroblastoma^39^ and more recently to function as a biomarker for tumor progression in Non-Small Cell Lung Cancer (NSCLS)^40^. MDK (FC = +2.82, p-adj = 0.0253) was previously identified as a neuroblastoma-secreted factor potentially involved in immune regulation in the TME of neuroblastoma^24^. To identify common biological processes that are mediated by secreted factors, we performed pathway analysis on the significantly (p-adj < 0.05) upregulated and downregulated proteins in the tumoroids compared to non-malignant controls. Processes associated with RNA splicing were significantly upregulated in the secretomes of tumoroids (top 5 proteins: RNA-binding motif protein (RBMX, FC = +2.73, p-adj = 0.0017), Heterogeneous nuclear ribonucleoproteins C1_C2 (HNRNPC, FC = 2.68, p-adj = 0.0033), Small nuclear ribonucleoprotein G (SNRPG, FC = 2.50, p-adj = 0.0065), RNA-splicing ligase RtcB homolog (RTCB, FC = 2.52, p-adj = 0.0067), Nucleolin (NCL, FC = 2.58, p-adj = 0.0075)) possibly a sign of a dysregulation of RNA splicing, an emerging hallmark of cancer^41^ (Fig. 2D).

In the top 100 most abundantly secreted proteins across tumoroids, we also identified High Mobility Group Box 1 (HMGB1), a well-known immunomodulatory protein in the TME of solid tumors^42^, as well as neuroblastoma-associated proteins Dopamine Beta-Hydroxylase (DBH) and Chromogranin B (CHGB)^43,44^ (Fig. 2E, Supplementary Fig. 2B). In addition, the well-known immunosuppressive factors TGFB1 and TGFB2 were identified in the secretomes of some of the tumoroids. Collectively, these provided strong support that our panel of neuroblastoma secretomes is representative of neuroblastomas *in vivo*, and reliable as a source to find physiologically relevant immunosuppressive factors.

Comparison of the MYCN non-amplified and MYCN amplified tumoroids identified three factors that were significantly more secreted by MYCN amplified tumoroids: NCL and Nucleoside Diphosphate Kinase 1/2 (NME1/2) (Supplementary Fig. 2C). These factors are highly associated with MYCN expression and prognostic for tumor progression in neuroblastoma, emphasizing the validity of our method^45,46^. Collectively, this suggests that MYCN activity can have far-reaching consequences not just in tumorigenesis, but perhaps also can influence the secretome and thereby affect immune cells in proximity of the tumor.

This dataset provides a deep characterization of the neuroblastoma secretome and its potential impact on shaping and interacting with the tumor microenvironment (TME). Next, to pinpoint factors specifically involved in immunosuppression, we compared the functional impacts of different secretomes on T cell activation.

### Different tumoroid secretomes exhibit varying degrees of T cell suppression

To determine the suppressive capacity of the tumoroid secretomes on T cell activation and cytotoxicity, healthy donor T cells were activated in the presence of 10µg/mL concentrated secretomes. After a 4 day incubation period, proliferation, activation and cytotoxicity markers were evaluated by flow cytometry (Fig. 3A, Supplementary Fig 3A). On average between all donors, 62.3% of the T cells proliferated without the addition of secretome (stimulated control). With the addition of tumoroid secretome, this percentage of proliferated T cells decreased, at most to 47.5%. This translated to 23.5% suppression when normalized to the stimulated control (Fig. 3B-C). Notably, virtually all tumoroids induced some degree, though distinct degrees, of T cell suppression. In contrast, the secretomes from both fibroblast, non-malignant controls (MaxFIB293 and MaxFIB272) promoted T cell proliferation. Both activation marker CD25 and cytotoxicity marker Granzyme B decreased in the presence of tumoroid secretomes (Fig. 3D-E). The suppression of CD25 expression reached a maximum of 38.8% (Fig. 3F) and the average suppression of Granzyme B ranged from 9.4 to 54.4% (Fig. 3G). The suppression rates of all three markers, i.e. proliferation, CD25 and Granzyme B, significantly correlated with each other across tumoroids indicating that the secretomes had a broad effect on T cell activity (Fig. 3H-J). Since we observed that some tumoroid secretomes induced high levels of suppression while others induced lower levels of suppression, we constructed a tumoroid-specific suppression score by normalizing and adding up the average tumoroid-specific suppression rates for proliferation, CD25 and Granzyme B. The secretome of AMC700T proved to be most suppressive and that of MaxNB400 the least suppressive (Fig. 3K).

**Figure 3:**
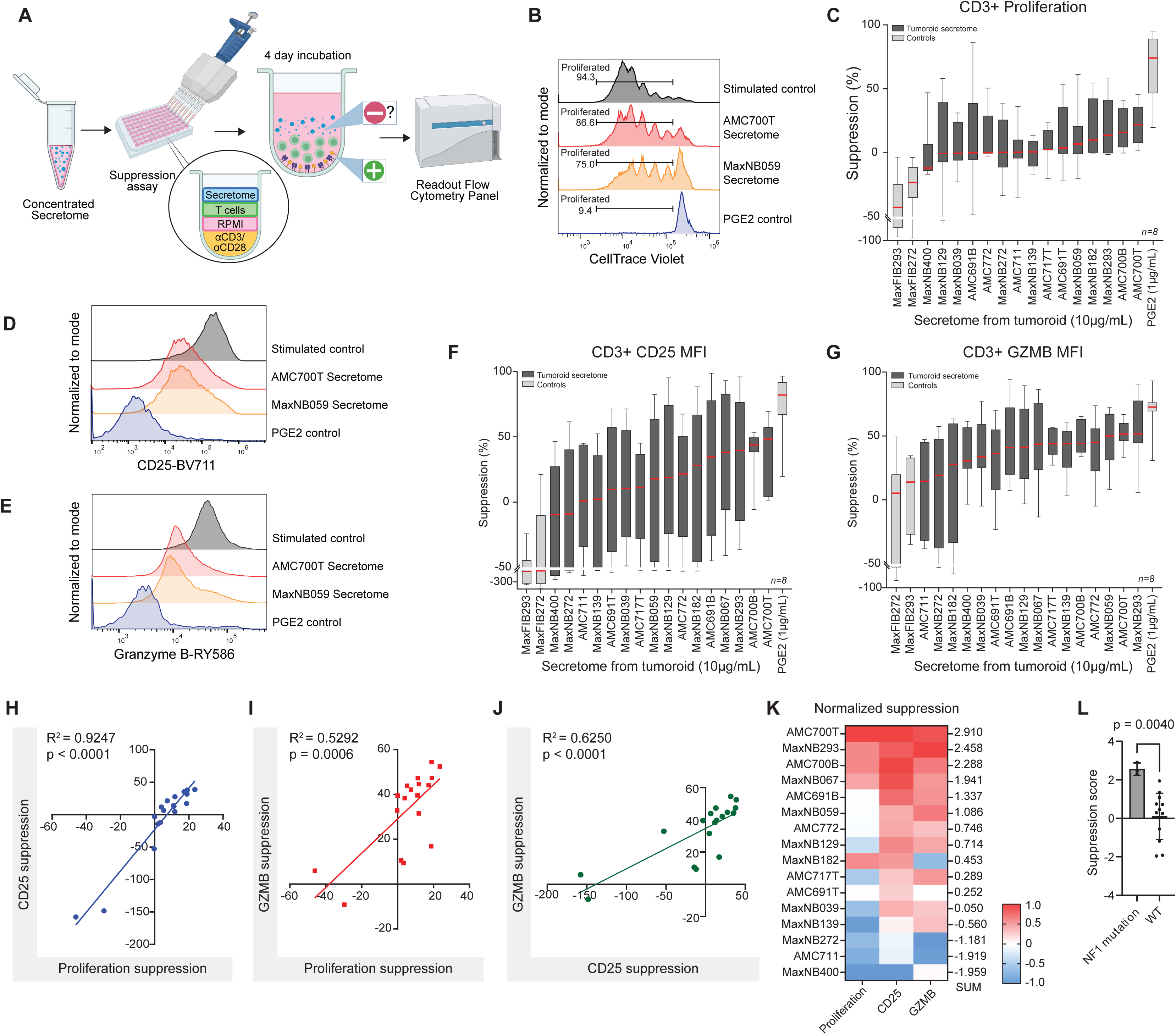
Determining suppression score of secretomes on T cells. A: Schematic representation of secretome suppression assay. **B:** Representative graph of T cell proliferation (gated as CellTrace Violet^(-)^ cells) showing the positive control (grey) and proliferation upon addition of secretome from AMC700T (red) and MaxNB059 (orange) or the Prostaglandin E2 (PGE2) control for suppression (blue). **C:** Bar graph showing the normalized suppression of T cell proliferation for each tumoroid secretome. The percentage of proliferated T cells was normalized to the stimulated control. The red line indicates the median. *N=8 healthy donors.* **D:** Histogram of CD25 expression on stimulated control (grey), secretome from AMC700T (red) and from MaxNB059 (orange) or the Prostaglandin E2 (PGE2) control for suppression (blue). **E:** Histogram of Granzyme B expression on stimulated control (grey), secretome from AMC700T (red) and from MaxNB059 (orange) or the Prostaglandin E2 (PGE2) control for suppression (blue). **F:** Normalized suppression of CD25 expression for each tumoroid secretome. The MFI of CD25 expression on T cells was normalized to the stimulated control. The red line indicates the median. *N=8 healthy donors.* **G:** Normalized suppression of Granzyme B expression for each tumoroid secretome. The MFI of Granzyme B expression on T cells was normalized to the stimulated control. The red line indicates the median. *N=8 healthy donors.* **H:** Correlation between suppression based on proliferation (x-axis) and CD25 (y-axis). Each dot represents the average suppression of secretome on *n=8 healthy donors.* **I:** Correlation between suppression based on proliferation (x-axis) and Granzyme B (y-axis). **J:** Correlation between suppression based on CD25 (x-axis) and Granzyme B (y-axis). **K:** Heatmap representing the min-max scaled suppression for each tumoroid’s secretome, with the sum of each parameter as suppression score. **L:** Comparison between suppression scores of NF1 mutated tumoroids and Wild Type (WT). Statistics show the unpaired t-test p-value.

Analysis of associations between genetic alterations of the tumoroids and the suppression scores revealed that tumors with an NF1 alteration (AMC700T and MaxNB293) had a significantly higher suppression score than non-NF1 altered tumors (Fig. 3L). NF1 is a tumor suppressor and when mutated in neuroblastoma can cause retinoic acid resistance as well as RAS-MEK signaling activation^47^. Cytokine levels in the TME of NF1-mutated tumors can be altered, but the direct effects of NF1-altered tumors on T cells are still unclear^48^. We did not observe an association with MYCN amplification, 1p loss, 11q loss or 17q gain with suppression (Supplementary Fig. 3B).

Taken together, we observed that neuroblastoma secretions can induce a spectrum of suppressive effects on T cells, which we translated into a suppression score associated with each neuroblastoma tumoroid line. This ranking system forms the basis to subsequently shortlist individual immunosuppressive factors (by correlation) to further narrow down to the specific source of T cell suppression.

### Neuropeptide Y highly correlates with secretome suppression scores

To determine which factors in the secretome drive T cell suppression, we compared the secretome contents as measured by LC-MS with the sum suppression scores determined in figure 3K. First, we compared the top 5 most suppressive tumoroid secretomes with the 5 least suppressive secretomes. 196 proteins were significantly (p < 0.05) upregulated in the most suppressive secretomes (Fig. 4A). Pathways upregulated in these 196 proteins included several pathways related to cell cycle progression (Orc1 removal from chromatin), translation, and Roundabout (ROBO) receptor signaling, which has been associated with migration, metastasis and immunosuppression (Supplementary Fig. 4A) ^49,50^. Next to that, we observed an upregulation of nervous system development or axon guidance pathways, underlining the neuronal features of neuroblastoma, possibly indicating a more malignant phenotype. To further narrow down potentially immunosuppressive factors, we assessed the correlation between the suppression score and all protein intensities measured by LC-MS. 56 unique proteins significantly positively correlated with the suppression scores (Fig. 4B). Among these, 29 were also significantly higher in the top 5 most immunosuppressive lines (Fig. 4C). 9 proteins were previously reported to correlate with immune cell infiltration within the TME (ACOT7, ATPC1A, CKMT1A, DDC, EWSR1, FAM19A5, HINT, MCM5, MAPT), but no direct effects on T cell activity or other immunomodulatory roles were described (Supplementary Table 2). Three additional factors – NCAM1, NMT, and NPY – were reported to directly affect immune cells within the TME, with NPY appearing to be most noteworthy as it was described to directly affect T cell proliferation^51^. Moreover, in neuroblastoma, NPY promotes tumor growth and vascularization and is particularly secreted by aggressive neuroblastomas with a poor prognosis (Supplementary Table 2)^51–53^. Based on these previous findings and the remarkably high association of NPY with T cell suppression scores in both statistical tests (Fig. 4D), we selected NPY for further investigation as a potential neuroblastoma-derived immunosuppressive factor.

**Figure 4:**
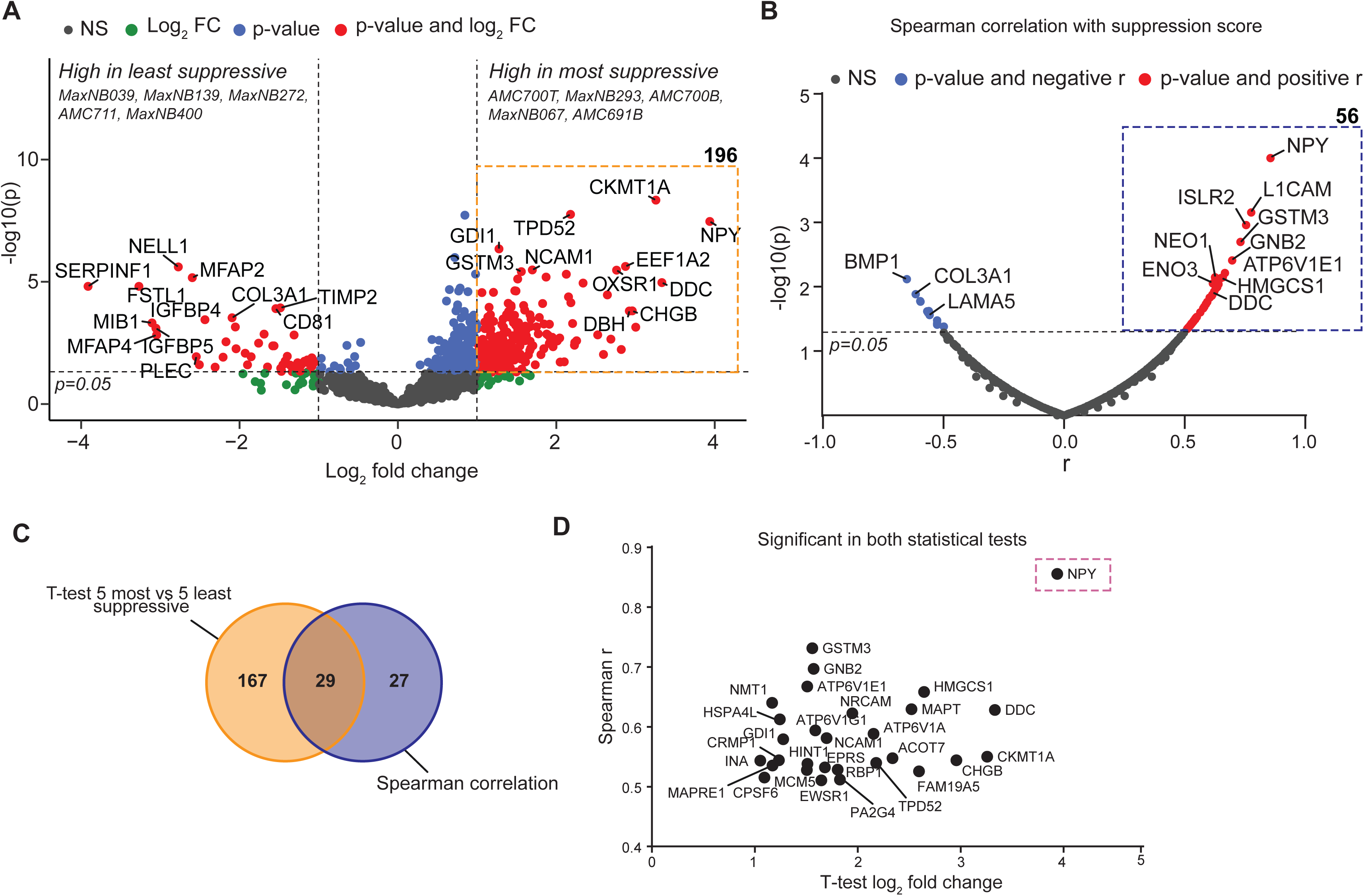
Using suppression score to analyze LC/MS data. A: Student’s t-test comparing the five most suppressive secretomes to the five least suppressive secretomes. Dashed orange box indicates proteins significantly upregulated in the most suppressive secretomes. **B:** Spearman correlation of secretome protein content measured by LC-MS with suppression score of each secretome. Dashed blue box indicates proteins with a significant positive correlation. **C:** Venn-diagram of proteins from Fig. 4A and Fig. 4B. **D:** XY-plot of the 29 proteins overlapping in Fig. 4C. Log2(FC) from Fig. 4A on x-axis and correlation (r) from Fig. 4B on y-axis. NPY has been annotated with a box.

### NPY does not cause direct suppression of T cell activation

To dissect possible mechanisms directly linking NPY to T cell activation, we performed extensive experiments involving (1) addition of recombinant human NPY protein to activated T cells, (2) treating activated T cells with small molecule NPY receptor antagonist, and (3) treating a functional neuroblastoma-targeting CAR-T cell with a NPY receptor antagonist. There are two active forms of NPY: the full-length (NPY-w) and the truncated NPY-3-36 ^54^. Due to the shotgun nature of proteomics, we could not distinguish the two isoforms in the secretome discovery data, but instead investigated the effect of both NPY active forms on direct T cell activation (Supplementary Fig. 5A). Direct addition of recombinant NPY to T cells did not significantly change proliferation, CD25 or Granzyme B expression, suggesting that NPY does not have a direct effect in regulating T cell activation (Supplementary figure 5B).

We next investigated if NPY may instead have autocrine effects on neuroblastoma tumoroids to induce an NPY-adapted secretome that is in turn suppressive on T cell activity. By ELISA, we quantified the abundance of secreted NPY across the full panel of tumoroid lines (Supplementary Fig. 5C), and proceeded with MaxNB400 (low suppression score, low NPY) for subsequent NPY supplementation experiments, and MaxNB293 (high suppression score, high NPY) for NPY blocking experiments, respectively. First, we assessed whether exposure of MaxNB400 to recombinant NPY could induce a more suppressive secretome (Supplementary Fig. 5D). Prior treatment of MaxNB400 tumoroids (low suppression score, low NPY) with recombinant NPY did not significantly alter the response of T cells that received the “NPY-adapted secretome” (Supplementary Fig. 5E). Blockade of NPY receptor on MaxNB293 tumoroids, which might abrogate autocrine signaling in MaxNB293 tumoroids (Supplementary Fig. 5F), similarly did not change the response of T cells that received the “NPY-R antagonist-adapted secretome” (Supplementary Fig. 5G). Collectively, these extensive data reveal that neuroblastoma tumoroids do not respond to autocrine NPY stimulation to alter their secretome to a more T cell suppressive composition.

Finally, we treated B7-H3 targeting CAR-T cells with an NPY-1 receptor antagonist in a co-culture with the highly suppressive MaxNB293 to evaluate the effect of NPY receptor blockade on the killing efficacy of CAR-T cells (Supplementary Fig. 5H)^51^. MaxNB293 did have B7-H3 expression and was effectively killed by the B7-H3 CAR-T cells, but no increased killing was observed upon NPY receptor blockade, suggesting a lack of a direct effect of NPY on (CAR) T cells (Supplementary Fig. 5I,J). In conclusion, we found no experimental evidence for a direct or indirect effect of NPY on T cell activation, though replicating TME interactions *in vitro* remains challenging.

### Immunosuppression is associated with extracellular vesicle secretion

Since NPY secretion was highly associated with immunosuppressive characteristics of neuroblastoma tumoroids, we next considered if NPY could be a biomarker of an immunosuppressive secretion phenotype to indicate poor outcome of T cell therapy, rather than exerting suppressive effects by itself. To evaluate this hypothesis, we reanalyzed the LC-MS data, to identify proteins which correlate with NPY secretion levels (Fig. 5A). 210 proteins were significantly and positively correlated with NPY secretion. Reactome pathway analysis highlighted pathways related to *Translation*, as well as *Membrane Trafficking* and *Vesicle-Mediated Transport*, suggesting that NPY may serve as an indicator of Extracellular Vesicle (EV) secretion (Fig. 5B).

**Figure 5:**
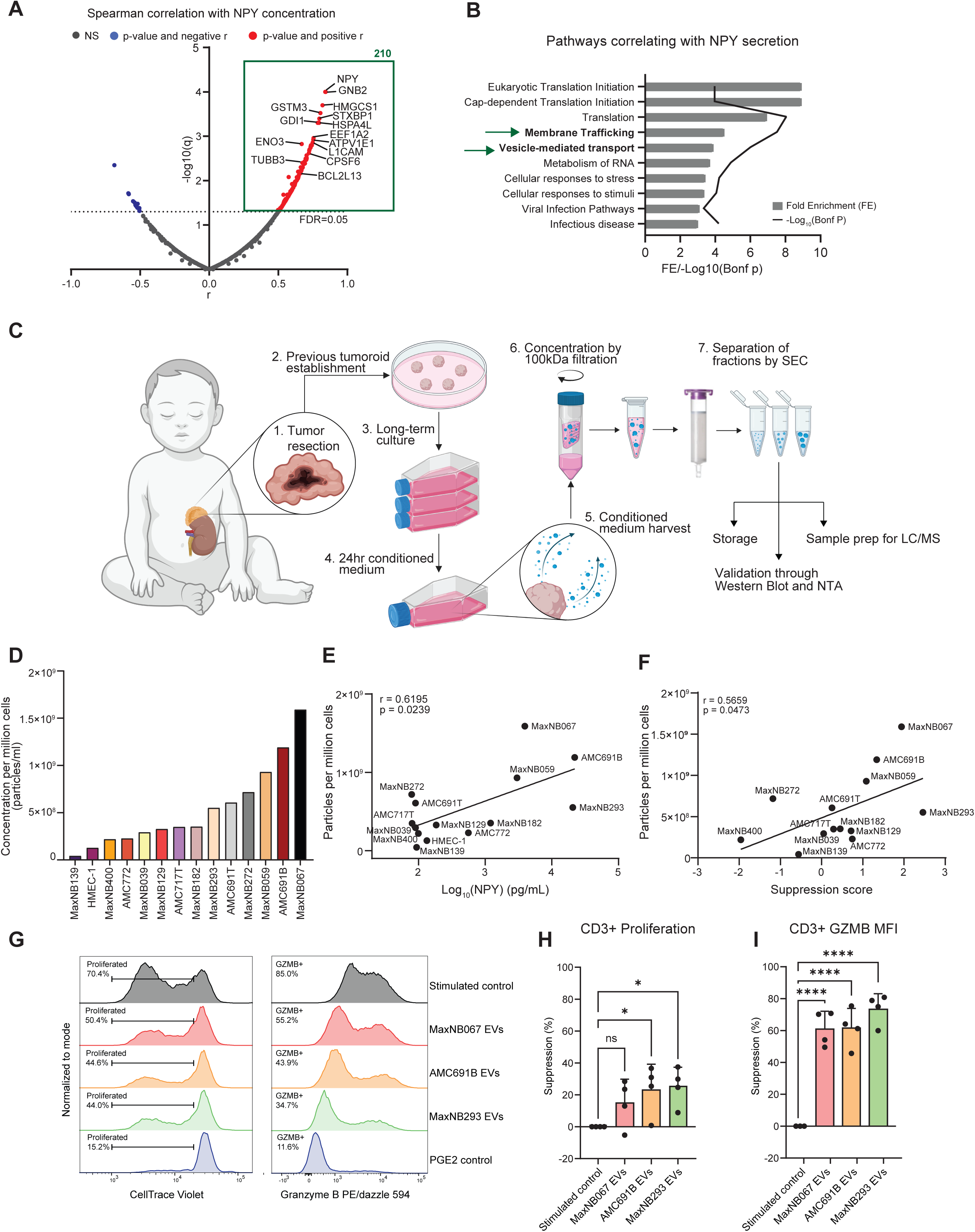
A: Spearman correlation between NPY secretion based on ELISA data from Supplementary Fig. 5C and all proteins in secretome data. **B:** Reactome pathway analysis with all 210 significant and positive correlation proteins from Fig. 5A. The top 10 pathways, ranking from the lowest Bonferroni p-values, with a Fold Enrichment above 3 and at least 10 proteins in the input, are shown. **C:** Schematic representation of protocol setup for isolation of EVs from the culture medium of neuroblastoma tumoroids. After SEC, fractions 5-9 were aliquoted for Western Blot and NTA analysis, LC-MS preparation and storage. **D:** Average concentration of particles/ml of EVs isolated from 14 tumoroid lines and HMEC-1, normalized to concentration per million cells of origin. **E:** XY-plot showing the correlation between NPY secretion as measure by ELISA and EV secretion as measured by particles/mL per million cells. **F:** XY-plot showing the correlation between suppression score (Fig. 3K) and EV secretion as measured by particles/mL per million cells. **G:** Representative graph of T cell proliferation (gated as CellTrace Violet^(-)^ cells) and Granzyme B expression showing the positive control (grey) and proliferation upon addition of EVs from MaxNB067 (red), AMC691B (orange) and MaxNB293 (green) or the Prostaglandin E2 (PGE2) control for suppression (blue). **H&I:** Bar graph showing the normalized suppression of T cell proliferation **(H)** and Granzyme B expression **(I)** by each tumoroid’s EVs. The percentage of proliferated T cells, c.q. MFI of granzyme B expression, was normalized to the stimulated control. *N=4 healthy donors.* Kruskall-Wallis with Dunn’s post hoc test.

To assess whether EV secretion could contribute T cell suppression, we isolated EVs from the conditioned media of 13 neuroblastoma tumoroids using size exclusion chromatography. We indeed observed that the number of EV particles secreted by each tumoroid correlated with secreted NPY levels (Fig. 5D,E), confirming that NPY is a biomarker for EV secretion. Moreover, the number of secreted EV particles significantly correlated with the tumoroid-specific suppression scores (Fig. 5F).

To subsequently test whether neuroblastoma-secreted EVs could suppress T cells *in vitro,* we activated T cells in the presence or absence of EVs, derived from the three most suppressive tumoroids (MaxNB067, AMC691B and MaxNB293). After 4 days, we observed that EVs from all three tumoroids lead to reduced proliferation and Granzyme B expression, and MaxNB293 EVs led to a trend of reduced CD25 expression, mirroring the previously obtained data from the secretome suppression assays in this study (Fig. 5G, H, I; Fig. 3C, F, G; Supplementary Fig 6 A, B). These data corroborate the concept that the EVs secreted by neuroblastoma tumoroids are the driving force behind the suppression of T cells.

## Discussion

CAR T-cell therapy for solid tumors is evolving to more effectively attack tumor cells by bypassing immunosuppressive effects in the TME. To do so, a detailed understanding of the immunosuppressive TME is pivotal. Here, we present a proteomic atlas of the neuroblastoma secretome generated with a unique library of 16 neuroblastoma patient-derived tumoroids. Using functional assays, we identified distinct suppressive effects of secretomes from different tumoroids on T cells, suggesting substantial inter-tumor heterogeneity in immunosuppressive capacity. Moreover, NPY expression proved an indicator for the rate of EV secretion. In turn, EVs secreted by neuroblastoma contribute to suppression of T cell proliferation and cytotoxicity in the TME.

Our analysis revealed that NPY is highly secreted by the most suppressive tumoroids, suggesting a potential association with immunosuppression in the TME. However, we could not confirm that NPY is the direct cause of immunosuppression, despite extensive and rigorous testing of receptor-ligand mechanisms and possible autocrine effects. NPY has previously been identified as a sympathetic neurotransmitter highly expressed by neuroblastomas^53^. Elevated levels of NPY in the serum of the neuroblastoma patients correlated with metastases, relapse and overall poor survival. Similarly, NPY was indicative of tumor progression, metastases and therapy resistance in other adult and pediatric cancers^55–58^. The correlation between NPY expression, immunosuppression and poor survival, supports the possibility that NPY may serve as a biomarker of severe disease, and a surrogate indicator of strong TME suppression of immune activation.

The complexity of tumor-induced immunosuppression became clearer with our finding that NPY secretion correlates with EV secretion, suggesting that EVs and their contents, rather than NPY itself, may drive immunosuppression. Tumor-derived EVs are increasingly studied for their role in immune modulation^59,60^. For example, EVs from nasopharyngeal carcinoma can recruit regulatory T cells, which suppresses anti-tumor responses, while prostate cancer-derived EVs can downregulate NKG2D on CD8^+^ T cells, impairing cytotoxicity^61,62^. EVs are also known to carry microRNAs (miRNAs) that can regulate target cell processes. In melanoma, tumor-derived EVs carry miRNAs which can activate the apoptosis pathway in CD4^+^ T cells in the TME or can induce the conversion of monocytes into myeloid derived suppressor cells, suppressing anti-tumor immunity^63,64^. In neuroblastoma, EVs have been shown to hinder CAR-T cell function *in vitro*, though the exact mechanism remains unknown^65^. In contrast, tumor-derived EVs displaying target antigens were recently shown to potentiate CAR T-cell activation and antitumor efficacy in neuroblastoma models^66^. Whether this benefit overcomes the immunosuppressive cargo which EVs may also carry in a physiologic setting, remains an open question.

A unique aspect of our study is our in-depth dataset of secretomes harvested from 16 patient-derived neuroblastoma tumoroids with varying genetic backgrounds, representing a valuable reference dataset for exploration of soluble factors released by neuroblastoma. While previous studies have investigated the secretome of neuroblastoma, most analyzed a small set of models, consisting of classical cell lines, which may be less representative for patient tumors^67,68^. In addition, we had access to two non-malignant lines derived from matched patient tumors. We used an unbiased approach to analyze the suppressive effects of the secretomes on T cell activation and revealed a role for EVs in T cell suppression. Our follow-up investigation will focus on proteomic profiling of the EVs isolated in this study to shed light on the exact mechanisms contributing to immunosuppression of T cells.

A limitation of our study is the lack of a complete TME to analyze the full complexity of soluble mediators interacting with and being derived from different cell types within the TME, such as macrophages or stromal cells, since they might contribute to the regulation of immune responses. Further validation of promising target proteins could therefore benefit from evaluation in the presence of a functional TME, such as in complex co-culture systems or *in vivo* studies.

From a translational perspective, several strategies are being explored to target EVs in cancer therapy^25^. One approach involves the inhibition of EV secretion by tumor cells; however, selective inhibition without affecting healthy cells remains a challenge. Kosaka *et al.* demonstrated that neutral sphingomyelinase 2 (nSMase2) is essential for exosome secretion by cancer cells, and its inhibition with GW4869 has been shown to reduce tumor progression in colorectal cancer models, albeit without reported effects on immune activity^69,70^. Alternatively, targeting immunosuppressive EV-associated molecules presents a promising strategy. For instance in breast cancer, vesicle-bound microRNA-122 (miR-122) suppresses glucose metabolism in target cells, facilitating the formation of a pre-metastatic niche and impairing cellular function within the TME^71^. Inhibition of miR-122 has been shown to restore glucose uptake in affected cells and reduce metastases. Further investigation into the immunosuppressive components of neuroblastoma-derived EVs is warranted to aid the development of targeted immunotherapeutic interventions.

In summary, we demonstrated that immunosuppressive neuroblastomas secrete high levels of EVs, reducing T cell functionality and impairing patient prognosis. With this, we hope to facilitate the further development of effective (immune)therapies for patients with neuroblastoma.

## Acknowledgements

This work received funding from Villa Joep (Joining forces to activate T cell immunity against High-Risk Neuroblastoma) and Veni (Release the beast: Boosting CAR-T cell immunotherapy for neuroblastoma, 09150162010022, which is partly financed by the Dutch Research Council (NWO) and by ZonMW). JS received travel grants from the Cultuurfonds and Academy Ter Meulen Grant. RB received travel grants from the Nijbakker-Morra foundation and Fonds Bemolt. J.A. received funding from the Great Ormond Street NIHR Biomedical Research centre and “Research in Childhood Cancer”. W.W. is supported by Singapore Immunology Network (SIgN), Agency for Science, Technology and Research (A*STAR); Biomedical Research Council (BMRC), Core Research Fund for use-inspired basic research (UIBR) and IAF-PP Project H22J2a0043, and Singapore National Medical Research Council (NMRC) project MOH-001401-00.

---

**Supplementary Figure 1.** :**A:** Oncoplot showing the genetic aberrations found in the panel of 16 neuroblastoma tumoroids as detected by WGS (snv = single nucleotide variant, LOH = Loss Of Heterozygosity). Also the treatment stage of the tumor samples is depicted. **B:** XY-plot showing the correlation between total protein concentration of concentrated secretome with cell count in one T175 flask per line. R^2^ and p-value depicted by simple linear regression analysis.

**Supplementary Figure 2:** A: Bar plot indication the proportion of unique proteins identified per secretome. Secretome or SignalP + Exocarta annotated proteins are annotated in blue. Non-secretome annotated are colored in yellow. **B:** Ranking of proteins in secretomes. iBAQ (intensity Based Absolute Quantification) indicates the protein’s non-normalized intensity divided by the number of measurable peptides and, hence, indicating the relative abundance of the protein in the sample. Vertical dotted line indicates rank 100. All factors mentioned in the main text are annotated. **C:** Volcano plot comparing all MYCN-amplified (A) secretomes to MYCN Non-Amplified (NA).

**Supplementary Figure 3:** A: Gating strategy for suppression assay read-out. **B:** Comparisons between groups in genetic alterations of tumoroids and their respective secretome’s suppression score. Statistics show the unpaired t-test p-value. WT=Wild Type.

**Supplementary Figure 4:** A: Reactome Pathway analysis for 196 proteins up in suppressive secretomes. Top 10 pathways with lowest Bonferroni P-value and a Fold Enrichment > 5 are shown.

**Supplementary Figure 5:** A: NPY peptide availability in LC-MS data of 7 tumoroid lines that had NPY in their secretome. Black squared at the bottom represent the Trypsin cleavage sites in the whole NPY peptide. **B:** Frequency of populations positive for the indicated activation markers within CD3^+^ T cells activated in the presence or absence of recombinant human NPY. Blue bars represent stimulated control, red bar the T cells treated with 0.01µM NPY-w and the green bar with 0.01µM NPY-3-36. *N=6 healthy T cell donors.* **C:** NPY ELISA of all secretomes. 500µg/mL total protein content for each secretome was loaded on the plates. Green triangle represents suppression score. Green arrows represent selected tumoroids for further testing. **D:** Schematic representation of the hypothesized effect of NPY treatment on MaxNB400 secretome. **E:** Frequency of populations positive for the indicated activation markers within CD3^+^ T cells activated in the presence of “baseline” MaxNB400 secretome or secretome from NPY-exposed MaxNB400, *n=6 healthy T cell donor*. **F:** Schematic representation of the hypothesized effect of NPY-receptor antagonists treatment on the MaxNB293 secretome. **G:** Frequency of population positive for the indicated activation markers within CD3^+^ T cells activated in the presence of “baseline” MaxNB293 secretome or secretome from MaxNB293 treated with NPY-receptor antagonists, *n=6 healthy T cell donor*. **H:** FACS staining for B7-H3 expression on MaxNB293. **I:** Schematic representation of α-NPY1R treatment on CAR T-cells to increase killing capacity. **J:** Caspase-3/7-green readout of killing assay in IncuCyte. B7-H3 targeting CAR-T cells were incubated with α-NPY1R (1µM) for 24 hours before the co-culture. 30.000 single cells of MaxNB293 were plated 48 hours before start of the co-culture and incubated to form back into spheres before treated CAR-T cells were added in a 1:10 Effector : Target ratio. Data were normalized to tumoroid only condition. *N=3 CAR T-cell donors*.

**Supplementary Figure 6:** A: Representative graph of CD25 expression showing the positive control (grey) and its reduction in activation upon addition of EVs from MaxNB067 (red), AMC691B (orange) and MaxNB293 (green) or the Prostaglandin E2 (PGE2) control for suppression (blue). **B:** Bar graph showing the normalized suppression of T cell activation by each tumoroid’s EVs. The MFI of CD25 expression, was normalized to the stimulated control. *N=4 healthy donors.* Kruskall-Wallis with Dunn’s post hoc test.

**Supplementary Table 1:** Overview of all tumoroids and controls used for secretome harvest. A picture was taken after 24hr conditioning of empty medium. The count of the cells after harvest per one full T175 and the protein content of the concentrated whole secretome is given.

**Supplementary Table 2:** Literature screening of 29 selected secreted factors

| Gene | Protein ID | Full name | Description | Known immunoregulatory function? | References |
| --- | --- | --- | --- | --- | --- |
| **ACOT7** | O00154 | Acyl-CoA thioesterase 7 | Role in cell cycle, proliferation and glucose metabolism. Overexpressed in tumor. | Involved in TME immune cell infiltration | 72 |
| **ATP6V1A** | P38606 | ATPase H<sup>+</sup> transporting V1 subunit A | Subunit of enzyme to regulate lysosomes and extracellular vesicles (EV). Is reduced in hypoxia, promoting EV secretion. | Positive correlation with immune infiltration in some malignancies (TIMER 2.0)<sup>73</sup> | 74,75 |
| **ATP6V1E1** | P36543 | ATPase H<sup>+</sup> transporting V1 subunit E1 | Subunit of enzyme to regulate lysosomes and extracellular vesicles (EV). | Not clearly found | 74,76 |
| **ATP6V1G1** | O75348 | ATPase H<sup>+</sup> Transporting V1 Subunit G1 | Subunit of enzyme to regulate lysosomes and extracellular vesicles (EV). | Not clearly found | 74 |
| **CHGB** | P05060 | Chromogranin B | Located in secretory granules of neuroendocrine cells; high in tumor; involved in processing of peptides in secretory granules | Involved in EMT induced by M2 macrophages in gliomas, and thus metastases | 77,78 |
| **CKMT1A** | P12532 | mitochondrial creatine kinase 1A | Expressed on mitochondrial surface and involved in transfer of phosphate groups | Negative association with CD8<sup>+</sup> infiltration in some tumors | 79 |
| **CPSF6** | Q16630 | Cleavage And Polyadenylation Specific Factor 6 | Involved in cancer cell proliferation | Not found | 80 |
| **CRMP1** | Q14194 | Collapsin response mediator protein 1 / Dihydropyrimidinase-related protein 1 | Located in cytosol exclusively in the nervous system / intracellular mediators of semaphorin 3A signaling pathway involved in axon growth and branching during neural development. Can suppress EMT in prostate cancer | Not found | 81 |
| **DDC** | P20711 | L-dopa decarboxylase / | Involved in biosynthesis of catecholamines | In clear cell renal cell carcinoma a positive correlation with a more immunosuppressive TME | 82,83 |
| **EPRS** | P07814 | glutamyl-prolyl-tRNA synthetase | Located in the cytoplasm; function within multisynthetase complex and catalyzes the attachment of L-glutamate and L-proline to their cognate tRNAs | Not found | 84 |
| **EWSR1** | Q01844 | Ewing Sarcoma RNA binding Protein 1 | Located in the nucleus / regulates transcription and RNA splicing / EWSR1::FLI1 initiate progression of Ewing sarcoma | Negative regulatory role for EWSR1 in constraining GC B cell responses (Expressed on B cells) | 85,86 |
| **FAM19A5** | Q7Z5A7 | Family With Sequence Similarity 19 / TAFA-5 | Secreted / highly expressed in the brain and adipose tissues / prominent in metabolic diseases | Not directly; S1PR2 (a FAM19A5 binder) is associated with immune infiltration (TIMER 2.0)<sup>73</sup> | 87,88 |
| **GDI1** | P31150 | guanine nucleotide dissociation inhibitor-1 | Located in cytoplasm / GDI1 binds to GDP-bound form of Rab proteins, keeping them inactive | Not found | 89 |
| **GNB2** | P62879 | G Protein Subunit Beta 2 | Located in cell membrane and cytoplasm / Subunit of Heterotrimeric G Protein | Not found | 90 |
| **GSTM3** | P21266 | Glutathione S-transferase Mu 3 | Located in cytoplasm / Detoxification of electrophilic compounds, such as cancer-causing toxins, anticarcinogens and products of oxidative stress via conjugating with glutathione | Not found | 91 |
| **HINT1** | P49773 | Adenosine 5'-monophosphoramidase HINT1 | Located in cytoplasm and nucleus / hydrolyzes purine nucleotide phosphoramidates substrates / Tumor suppressor gene | High expression of HINT1 was associated with the decreased infiltration of cluster of differentiated CD4+ T cells (TIMER 2.0)<sup>73</sup> | 92,93 |
| **HMGCS1** | Q01581 | 3-hydroxy-3-methylglutaryl-CoA synthase 1 | Located in cytoplasm / Catalyses conversion of acetoacetyl-CoA to(HMG-CoA) / Control mevalonate pathway (synthesis of cholesterol) | Indirect: inhibition of the mevalonate pathway promotes pyroptosis, which induces immune cell infiltration through inflammatory cytokines | 94 |
| **HSPA4L** | O95757 | Heat shock 70 kDa protein 4L | Heat Shock proteins can protect the cell against the harmful effect of protein aggregation after a heat shock. High expression in leukaemia | HSPA4 (so not L) has been written about in the context of immune regulation in hepatocellular carcinoma | 95,96 |
| **INA** | Q16352 | Internexin Neuronal Intermediate Filament Protein Alpha | Neurofilament; part of the exoskeleton of neurons | Not found | 97,98 |
| **MAPRE1** | Q15691 | Microtubule associated protein RP/EB family member 1 | Localizes to the microtubules; during mitosis of the cells associated with the centrosomes and spindle microtubules. Possible association with chromosomal instability. | Not found | 99,100 |
| **MAPT** | P10636 | Microtubule-associated protein tau | Mostly found in neurons, where it stabilizes microtubules | Correlation with lower immune infiltration | 101 |
| **MCM5** | P33992 | Minichromosome maintenance complex component 5 | Transcription factor, which forms a complex involved in DNA replication; interaction with STAT1 | Positive correlation with immune infiltration | 102,103 |
| **NCAM1** | P13591 | Neural Cell Adhesion Molecule 1 | Cell-cell adhesion molecule expressed by many cancers; also known as CD56 expressed by NK and other immune cells | Yes, expressed by NK cells and can enhance NK cells | 104,105 |
| **NMT1** | P30419 | N-myristoyltransferase 1 | Enzyme catalysing in a process called myristylation for lipid modifications of proteins. Overexpressed in tumors and already discovered as an interesting target for cancer therapy. | Yes, plays a role in growth and differentiation in cells of leukocytic origin | 106,107 |
| **NPY** | P01303 | Neuropeptide Y | NPY is a transmitter between the nervous system and the immune system. | Suppresses T cell proliferation; induces Th2 polarization and suppresses NK activity; can also polarize macrophages to more M2 | 51,108,109 |
| **NRCAM** | Q92823 | Neuronal cell adhesion molecule | Adhesion molecule / neuronal receptor closely related to NCAM; has a role in neural development. | No; other related genes have functions in f.i. MS | 110 |
| **PA2G4** | Q9UQ80 | proliferation-associated 2G4 | Protein associated with cancer regulating cellular growth and differentiation. Role depends on the isoform and context. Associated with MYCN expression in neuroblastoma | Not clearly, there is a correlation with lower immune infiltration | 111 |
| **RBP1** | P09455 | Retinol binding protein 1 | Involved in the transportation of retinol; involved in several cancers | Not clearly; co-expression with immune checkpoints genes in several tumors | 112 |
| **TPD52** | P55327 | Tumor protein D52 | Protein highly overexpressed in tumor; In lung carcinoma there is a correlation with TPD52(L2) expression and the infiltration of immunosuppressive cells in the TME | Not clearly on T/NK. Some correlation with immunosuppressive TME; Seems to be expressed by plasma B cells | 113 |

## References

1. Korman, A. J., Garrett-Thomson, S. C. & Lonberg, N. The foundations of immune checkpoint blockade and the ipilimumab approval decennial. Nat Rev Drug Discov 21, 509–528 (2022).

2. Galluzzi, L., Chan, T. A., Kroemer, G., Wolchok, J. D. & López-Soto, A. The hallmarks of successful anticancer immunotherapy. Sci Transl Med 10, (2018).

3. June, C. H. & Sadelain, M. Chimeric Antigen Receptor Therapy. New England Journal of Medicine 379, 64–73 (2018).

4. Maude, S. L. et al. Tisagenlecleucel in Children and Young Adults with B-Cell Lymphoblastic Leukemia. New England Journal of Medicine 378, 439–448 (2018).

5. Hou, A. J., Chen, L. C. & Chen, Y. Y. Navigating CAR-T cells through the solid-tumour microenvironment. Nat Rev Drug Discov 20, 531–550 (2021).

6. Matthay, K. K. et al. Neuroblastoma. Nat Rev Dis Primers 2, 16078 (2016).

7. Matthay, K. K. et al. Advances in Risk Classification and Treatment Strategies for Neuroblastoma. Journal of Clinical Oncology 33, 3008–3017 (2015).

8. Cheung, N. K. V & Dyer, M. A. Neuroblastoma: Developmental biology, cancer genomics and immunotherapy. Nat Rev Cancer 13, 397–411 (2013).

9. Louis, C. U. et al. Antitumor activity and long-term fate of chimeric antigen receptor-positive T cells in patients with neuroblastoma. Blood 118, 6050–6 (2011).

10. Heczey, A. et al. CAR T Cells Administered in Combination with Lymphodepletion and PD-1 Inhibition to Patients with Neuroblastoma. Mol Ther 25, 2214–2224 (2017).

11. Zappa, E. et al. Adoptive cell therapy in paediatric extracranial solid tumours: current approaches and future challenges. Eur J Cancer 194, 113347 (2023).

12. Del Bufalo, F. et al. GD2-CART01 for Relapsed or Refractory High-Risk Neuroblastoma. N Engl J Med 388, 1284–1295 (2023).

13. Padgaonkar, M., Shendre, S., Chatterjee, P. & Banerjee, S. Cancer secretome: finding out hidden messages in extracellular secretions. Clinical and Translational Oncology 25, 1145–1155 (2023).

14. Doyle, L. & Wang, M. Overview of Extracellular Vesicles, Their Origin, Composition, Purpose, and Methods for Exosome Isolation and Analysis. Cells 8, 727 (2019).

15. Buzas, E. I. The roles of extracellular vesicles in the immune system. Nat Rev Immunol 0123456789, (2022).

16. Källberg, J. et al. Intratumor heterogeneity and cell secretome promote chemotherapy resistance and progression of colorectal cancer. Cell Death Dis 14, (2023).

17. Feizi, A., Banaei-Esfahani, A. & Nielsen, J. HCSD: The human cancer secretome database. Database (2015).

18. Ritchie, S., Reed, D. A., Pereira, B. A. & Timpson, P. The cancer cell secretome drives cooperative manipulation of the tumour microenvironment to accelerate tumourigenesis. Fac Rev 10, (2021).

19. Papaleo, E., Gromova, I. & Gromov, P. Gaining insights into cancer biology through exploration of the cancer secretome using proteomic and bioinformatic tools. Expert Rev Proteomics 14, 1021–1035 (2017).

20. Zeng, X. et al. Quantitative secretome analysis reveals the interactions between epithelia and tumor cells by in vitro modulating colon cancer microenvironment. J Proteomics 89, 51–70 (2013).

21. Massagué, J. TGFβ in Cancer. *Cell* **134**, 215–230 (2008).

22. Thomas, D. A. & Massagué, J. TGF-β directly targets cytotoxic T cell functions during tumor evasion of immune surveillance. Cancer Cell 8, 369–380 (2005).

23. Ciardiello, D., Elez, E., Tabernero, J. & Seoane, J. Clinical development of therapies targeting TGFβ: current knowledge and future perspectives. Annals of Oncology 31, 1336–1349 (2020).

24. Strijker, J. G. M. et al. Blocking MIF secretion enhances CAR T-cell efficacy against neuroblastoma. Eur J Cancer 218, 115263 (2025).

25. Wang, S. E. Extracellular vesicles in cancer therapy. Semin Cancer Biol 86, 296–309 (2022).

26. Ho, P.-C. & Liu, P.-S. Metabolic communication in tumors: a new layer of immunoregulation for immune evasion. J Immunother Cancer 4, 4 (2016).

27. Brand, A. et al. LDHA-Associated Lactic Acid Production Blunts Tumor Immunosurveillance by T and NK Cells. Cell Metab 24, 657–671 (2016).

28. Anderson, K. G., Stromnes, I. M. & Greenberg, P. D. Obstacles Posed by the Tumor Microenvironment to T cell Activity: A Case for Synergistic Therapies. Cancer Cell 31, 311–325 (2017).

29. Bate-Eya, L. T. et al. Newly-derived neuroblastoma cell lines propagated in serum-free media recapitulate the genotype and phenotype of primary neuroblastoma tumours. Eur J Cancer 50, 628–637 (2014).

30. Langenberg, K. P. S. et al. Implementation of paediatric precision oncology into clinical practice: The Individualized Therapies for Children with cancer program ‘iTHER’. Eur J Cancer 175, 311–325 (2022).

31. Langenberg, K. P. S. et al. Exploring high-throughput drug sensitivity testing in neuroblastoma cell lines and patient-derived tumor organoids in the era of precision medicine. Eur J Cancer 218, 115275 (2025).

32. Birley, K. et al. A novel anti-B7-H3 chimeric antigen receptor from a single-chain antibody library for immunotherapy of solid cancers. Mol Ther Oncolytics 26, 429–443 (2022).

33. Langenberg, K. P. S. et al. Exploring high-throughput drug sensitivity testing in neuroblastoma cell lines and patient-derived tumor organoids in the era of precision medicine. Eur J Cancer 218, (2025).

34. Ades, E. W. et al. HMEC-1: establishment of an immortalized human microvascular endothelial cell line. J Invest Dermatol 99, 683–90 (1992).

35. Uhlén, M. et al. The human secretome. *Sci Signal* **12**, (2019).

36. Mathivanan, S., Fahner, C. J., Reid, G. E. & Simpson, R. J. ExoCarta 2012: database of exosomal proteins, RNA and lipids. Nucleic Acids Res 40, D1241–D1244 (2012).

37. The human protein atlas. The Secretome. https://www.proteinatlas.org/humanproteome/tissue/secretome (2022).

38. Fhu, C. W. & Ali, A. Fatty Acid Synthase: An Emerging Target in Cancer. Molecules 25, 3935 (2020).

39. Wittke, I., et al. Neuroblastoma-Derived Sulfhydryl Oxidase, a New Member of the Sulfhydryl Oxidase/Quiescin6 Family, Regulates Sensitization to Interferon-Induced Cell Death in Human Neuroblastoma Cells. http://publication.celera.com. (2003).

40. Li, Y. et al. QSOX2 Is an E2F1 Target Gene and a Novel Serum Biomarker for Monitoring Tumor Growth and Predicting Survival in Advanced NSCLC. Front Cell Dev Biol 9, (2021).

41. Bradley, R. K. & Anczuków, O. RNA splicing dysregulation and the hallmarks of cancer. Nat Rev Cancer 23, 135–155 (2023).

42. Wen, J. et al. Knockdown of HMGB1 inhibits the crosstalk between oral squamous cell carcinoma cells and tumor-associated macrophages. Int Immunopharmacol 119, 110259 (2023).

43. Giudici, A. M., Sher, E., Pelagi, M., Clementi, F. & Zanini, A. Immunolocalization of secretogranin II, chromogranin A, and chromogranin B in differentiating human neuroblastoma cells. Eur J Cell Biol 58, 383–9 (1992).

44. van Zogchel, L. M. J. et al. Specific and Sensitive Detection of Neuroblastoma mRNA Markers by Multiplex RT-qPCR. Cancers (Basel) 13, 150 (2021).

45. Cangelosi, D. et al. Nucleolin expression has prognostic value in neuroblastoma patients. EBioMedicine 85, 104300 (2022).

46. Chicco, D., Sanavia, T. & Jurman, G. Signature literature review reveals AHCY, DPYSL3, and NME1 as the most recurrent prognostic genes for neuroblastoma. BioData Min 16, 7 (2023).

47. Hölzel, M. et al. NF1 Is a Tumor Suppressor in Neuroblastoma that Determines Retinoic Acid Response and Disease Outcome. Cell 142, 218–229 (2010).

48. Karmakar, S. & Reilly, K. M. The role of the immune system in neurofibromatosis type 1-associated nervous system tumors. CNS Oncol 6, 45–60 (2017).

49. Geraldo, L. H. et al. SLIT2/ROBO signaling in tumor-associated microglia and macrophages drives glioblastoma immunosuppression and vascular dysmorphia. Journal of Clinical Investigation 131, (2021).

50. Blockus, H. & Chédotal, A. Slit-Robo signaling. Development 143, 3037–3044 (2016).

51. Wheway, J., Herzog, H. & Mackay, F. The Y1 receptor for NPY: A key modulator of the adaptive immune system. Peptides (N.Y.) 28, 453–458 (2007).

52. Kitlinska, J. Neuropeptide Y (NPY) in neuroblastoma: Effect on growth and vascularization. Peptides (N.Y.) 28, 405–412 (2007).

53. Galli, S. et al. Neuropeptide Y as a Biomarker and Therapeutic Target for Neuroblastoma. American Journal of Pathology 186, 3040–3053 (2016).

54. Chen, W. C. et al. Neuropeptide Y Is an Immunomodulatory Factor: Direct and Indirect. Front Immunol 11, (2020).

55. Hong, S.-H. et al. High neuropeptide Y release associates with Ewing sarcoma bone dissemination - *in vivo* model of site-specific metastases. Oncotarget 6, 7151–7165 (2015).

56. Tilan, J. & Kitlinska, J. Neuropeptide Y (NPY) in tumor growth and progression: Lessons learned from pediatric oncology. Neuropeptides 55, 55–66 (2016).

57. Raunkilde, L. et al. NPY Gene Methylation in Circulating Tumor DNA as an Early Biomarker for Treatment Effect in Metastatic Colorectal Cancer. Cancers (Basel) 14, 4459 (2022).

58. Ding, Y. et al. Neuropeptide Y nerve paracrine regulation of prostate cancer oncogenesis and therapy resistance. Prostate 81, 58–71 (2021).

59. Mir, R. et al. Unlocking the Secrets of Extracellular Vesicles: Orchestrating Tumor Microenvironment Dynamics in Metastasis, Drug Resistance, and Immune Evasion. J Cancer 15, 6383–6415 (2024).

60. Reale, A., Khong, T. & Spencer, A. Extracellular Vesicles and Their Roles in the Tumor Immune Microenvironment. J Clin Med 11, 6892 (2022).

61. Mrizak, D. et al. Effect of Nasopharyngeal Carcinoma-Derived Exosomes on Human Regulatory T Cells. JNCI: Journal of the National Cancer Institute 107, (2015).

62. Lundholm, M. et al. Prostate Tumor-Derived Exosomes Down-Regulate NKG2D Expression on Natural Killer Cells and CD8+ T Cells: Mechanism of Immune Evasion. PLoS One 9, e108925 (2014).

63. Zhou, J. et al. Melanoma-released exosomes directly activate the mitochondrial apoptotic pathway of CD4+ T cells through their microRNA cargo. Exp Cell Res 371, 364–371 (2018).

64. Huber, V. et al. Tumor-derived microRNAs induce myeloid suppressor cells and predict immunotherapy resistance in melanoma. Journal of Clinical Investigation 128, 5505–5516 (2018).

65. Ali, S. et al. Tumor-Derived Extracellular Vesicles Impair CD171-Specific CD4+ CAR T Cell Efficacy. Front Immunol 11, (2020).

66. Giudice, A. M., et al. Target Antigen-Displaying Extracellular Vesicles Boost CAR T Cell Efficacy in Cell and Mouse Models of Neuroblastoma. Sci. Transl. Med vol. 17 https://www.science.org (2025).

67. Rozek, W., Kwasnik, M., Debski, J. & Zmudzinski, J. F. Mass spectrometry identification of granins and other proteins secreted by neuroblastoma cells. Tumor Biology 34, 1773–1781 (2013).

68. Zhang, H.-F. et al. A MYCN-independent mechanism mediating secretome reprogramming and metastasis in *MYCN* -amplified neuroblastoma. Sci Adv 9, (2023).

69. Kosaka, N. et al. Secretory Mechanisms and Intercellular Transfer of MicroRNAs in Living Cells. Journal of Biological Chemistry 285, 17442–17452 (2010).

70. Kosaka, N. et al. Neutral Sphingomyelinase 2 (nSMase2)-dependent Exosomal Transfer of Angiogenic MicroRNAs Regulate Cancer Cell Metastasis. Journal of Biological Chemistry 288, 10849–10859 (2013).

71. Fong, M. Y. et al. Breast-cancer-secreted miR-122 reprograms glucose metabolism in premetastatic niche to promote metastasis. Nat Cell Biol 17, 183–194 (2015).

72. Zheng, C. et al. Pan-Cancer Analysis and Experimental Validation Identify ACOT7 as a Novel Oncogene and Potential Therapeutic Target in Lung Adenocarcinoma. Cancers (Basel) 14, 4522 (2022).

73. Li, T. et al. TIMER2.0 for analysis of tumor-infiltrating immune cells. Nucleic Acids Res 48, W509–W514 (2020).

74. Li, X. et al. Comprehensive Analysis of ATP6V1s Family Members in Renal Clear Cell Carcinoma With Prognostic Values. Front Oncol 10, (2020).

75. Wang, X. et al. Hypoxia promotes EV secretion by impairing lysosomal homeostasis in HNSCC through negative regulation of ATP6V1A by HIF-1α. J Extracell Vesicles 12, (2023).

76. Feng, J. et al. Construction and validation of a novel cuproptosis-mitochondrion prognostic model related with tumor immunity in osteosarcoma. PLoS One 18, e0288180 (2023).

77. Yadav, G. P. et al. Chromogranin B (CHGB) is dimorphic and responsible for dominant anion channels delivered to cell surface via regulated secretion. Front Mol Neurosci 16, (2023).

78. Feng, P., Liu, S., Yuan, G. & Pan, Y. Association of M2 macrophages with EMT in glioma identified through combination of multi-omics and machine learning. Heliyon 10, e34119 (2024).

79. Yang, M., Liu, S., Xiong, Y., Zhao, J. & Deng, W. An integrative pan-cancer analysis of molecular characteristics and oncogenic role of mitochondrial creatine kinase 1A (CKMT1A) in human tumors. Sci Rep 12, 10025 (2022).

80. Liu, S. et al. CPSF6 regulates alternative polyadenylation and proliferation of cancer cells through phase separation. Cell Rep 42, 113197 (2023).

81. Cai, G. et al. Collapsin response mediator protein-1 (CRMP1) acts as an invasion and metastasis suppressor of prostate cancer via its suppression of epithelial–mesenchymal transition and remodeling of actin cytoskeleton organization. Oncogene 36, 546–558 (2017).

82. Kuhar, M., Couceyro, P. & Lambert, P. Basic Neurochemistry: Molecular, Cellular and Medical Aspects. in Basic Neurochemistry*, 6th edition* (eds. Siegel, G., Agranoff, B., Wayne Albers, R., Fisher, S. & Uhler, M.) (Lippincott-Raven, Philadelphia, 1999).

83. Chang, K. et al. Multi-omics profiles refine L-dopa decarboxylase (DDC) as a reliable biomarker for prognosis and immune microenvironment of clear cell renal cell carcinoma. Front Oncol 12, (2022).

84. Lee, E.-Y., Hwang, J. & Kim, M. H. Phosphocode-dependent glutamyl-prolyl-tRNA synthetase 1 signaling in immunity, metabolism, and disease. Exp Mol Med 55, 2116–2126 (2023).

85. Wang, Y. et al. Gammaherpesvirus-mediated repression reveals EWSR1 to be a negative regulator of B cell responses. Proceedings of the National Academy of Sciences 119, (2022).

86. Lee, J. et al. EWSR1, a multifunctional protein, regulates cellular function and aging via genetic and epigenetic pathways. Biochimica et Biophysica Acta (BBA) - Molecular Basis of Disease 1865, 1938–1945 (2019).

87. Park, M. Y. et al. FAM19A5, a brain-specific chemokine, inhibits RANKL-induced osteoclast formation through formyl peptide receptor 2. Sci Rep 7, 15575 (2017).

88. Cheng, J.-X. & Yu, K. New Discovered Adipokines Associated with the Pathogenesis of Obesity and Type 2 Diabetes. Diabetes Metab Syndr Obes **Volume** 15, 2381–2389 (2022).

89. Xie, X. et al. Overexpression of GDP dissociation inhibitor 1 gene associates with the invasiveness and poor outcomes of colorectal cancer. Bioengineered 12, 5595–5606 (2021).

90. Zhang, L. et al. A detailed multi-omics analysis of GNB2 gene in human cancers. Brazilian Journal of Biology 84, (2024).

91. Zhang, J. et al. Comprehensive analysis of the glutathione S-transferase Mu (GSTM) gene family in ovarian cancer identifies prognostic and expression significance. Front Oncol 12, (2022).

92. Dillenburg, M., Smith, J. & Wagner, C. R. The Many Faces of Histidine Triad Nucleotide Binding Protein 1 (HINT1). ACS Pharmacol Transl Sci 6, 1310–1322 (2023).

93. Wang, X., Zhou, M. & Jiang, L. The oncogenic and immunological roles of histidine triad nucleotide-binding protein 1 in human cancers and their experimental validation in the MCF-7 cell line. Ann Transl Med 11, 147–147 (2023).

94. Zhou, W. et al. Targeting the mevalonate pathway suppresses ARID1A-inactivated cancers by promoting pyroptosis. Cancer Cell 41, 740–756.e10 (2023).

95. Shang, B.-B., Chen, J., Wang, Z.-G. & Liu, H. Significant correlation between HSPA4 and prognosis and immune regulation in hepatocellular carcinoma. PeerJ 9, e12315 (2021).

96. Takahashi, H. et al. Identification of an overexpressed gene, HSPA4L, the product of which can provoke prevalent humoral immune responses in leukemia patients. Exp Hematol 35, 1091–1099 (2007).

97. Shea, T. B. & Beermann, M. L. Neuronal intermediate filament protein alpha-internexin facilitates axonal neurite elongation in neuroblastoma cells. Cell Motil Cytoskeleton 43, 322–33 (1999).

98. Yuan, A. et al. α-Internexin Is Structurally and Functionally Associated with the Neurofilament Triplet Proteins in the Mature CNS. The Journal of Neuroscience 26, 10006–10019 (2006).

99. Liang, X. et al. MAPRE1 promotes cell cycle progression of hepatocellular carcinoma cells by interacting with CDK2. Cell Biol Int 44, 2326–2333 (2020).

100. Shao, M., Li, W., Wang, S. & Liu, Z. Identification of key genes and pathways associated with esophageal squamous cell carcinoma development based on weighted gene correlation network analysis. J Cancer 11, 1393–1402 (2020).

101. Liu, Z. et al. A Novel Aging-Related Prognostic lncRNA Signature Correlated with Immune Cell Infiltration and Response to Immunotherapy in Breast Cancer. Molecules 28, 3283 (2023).

102. Sun, M. et al. The High Expression of Minichromosome Maintenance Complex Component 5 Is an Adverse Prognostic Factor in Lung Adenocarcinoma. Biomed Res Int 2022, 1–17 (2022).

103. Snyder, M., He, W. & Zhang, J. J. The DNA replication factor MCM5 is essential for Stat1-mediated transcriptional activation. Proceedings of the National Academy of Sciences 102, 14539–14544 (2005).

104. Picard, L. K., Claus, M., Fasbender, F. & Watzl, C. Human NK cells responses are enhanced by CD56 engagement. Eur J Immunol 52, 1441–1451 (2022).

105. Weledji, E. P. & Assob, J. C. The ubiquitous neural cell adhesion molecule (N-CAM). Annals of Medicine & Surgery 3, 77–81 (2014).

106. Gamma, J. et al. Validation of *NMT1* and *NMT2* As Novel Drug Targets in Adult Acute Myeloid Leukemia: Rationale for N-Myristoyltransferase Inhibition with Pclx-001 for Clinical Trials. Blood 138, 4344–4344 (2021).

107. Kumar, S., Singh, B., Dimmock, J. R. & Sharma, R. K. N-myristoyltransferase in the leukocytic development processes. Cell Tissue Res 345, 203–211 (2011).

108. Dimitrijević, M. & Stanojević, S. The intriguing mission of neuropeptide y in the immune system. Amino Acids 45, 41–53 (2013).

109. Profumo, E. et al. Neuropeptide Y Promotes Human M2 Macrophage Polarization and Enhances p62/SQSTM1-Dependent Autophagy and NRF2 Activation. Int J Mol Sci 23, (2022).

110. Conacci-Sorrell, M. E. et al. Nr-CAM is a target gene of the β-catenin/LEF-1 pathway in melanoma and colon cancer and its expression enhances motility and confers tumorigenesis. Genes Dev 16, 2058–2072 (2002).

111. Stevenson, B. W. et al. A structural view of PA2G4 isoforms with opposing functions in cancer. Journal of Biological Chemistry 295, 16100–16112 (2020).

112. Gao, L. et al. The RBP1–CKAP4 axis activates oncogenic autophagy and promotes cancer progression in oral squamous cell carcinoma. Cell Death Dis 11, 488 (2020).

113. Sruthi, K. K., Natani, S. & Ummanni, R. Tumor protein D52 (isoform 3) induces NF-κB – STAT3 mediated EMT driving neuroendocrine differentiation of prostate cancer cells. Int J Biochem Cell Biol 166, 106493 (2024).

