## Supplementary figures and table for "Proteomic profiling of the neuroblastoma secretome identifies extracellular vesicles as drivers of T cell suppression"

Supplementary figure 1

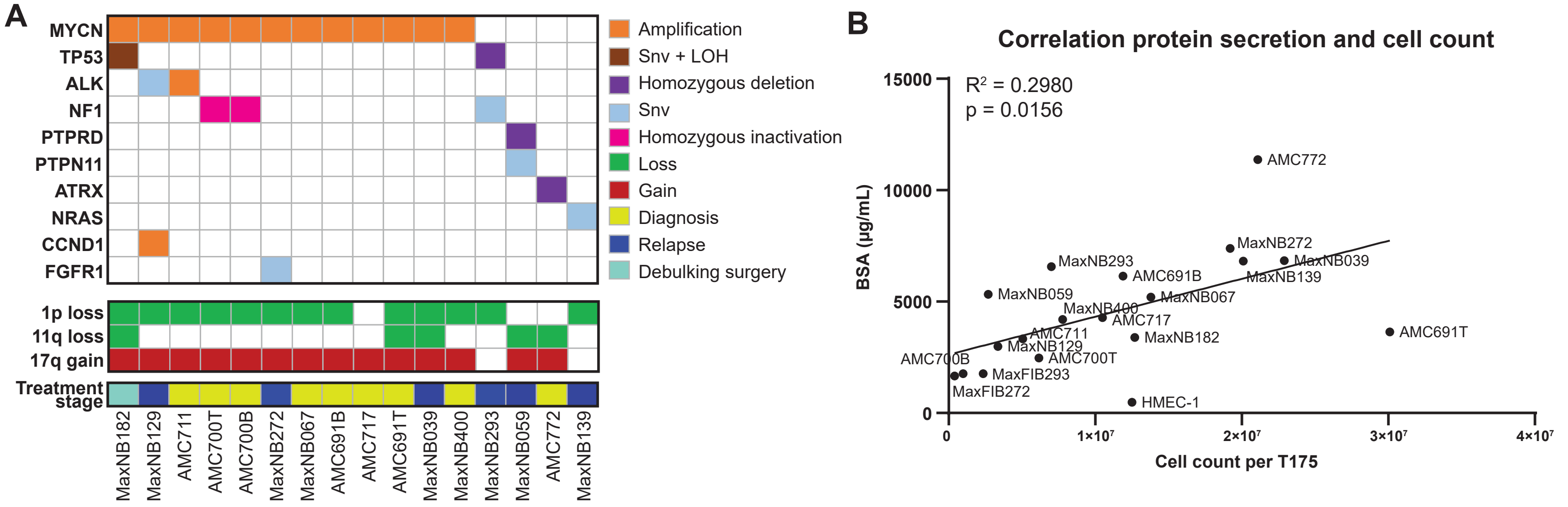

Supplementary Figure 2

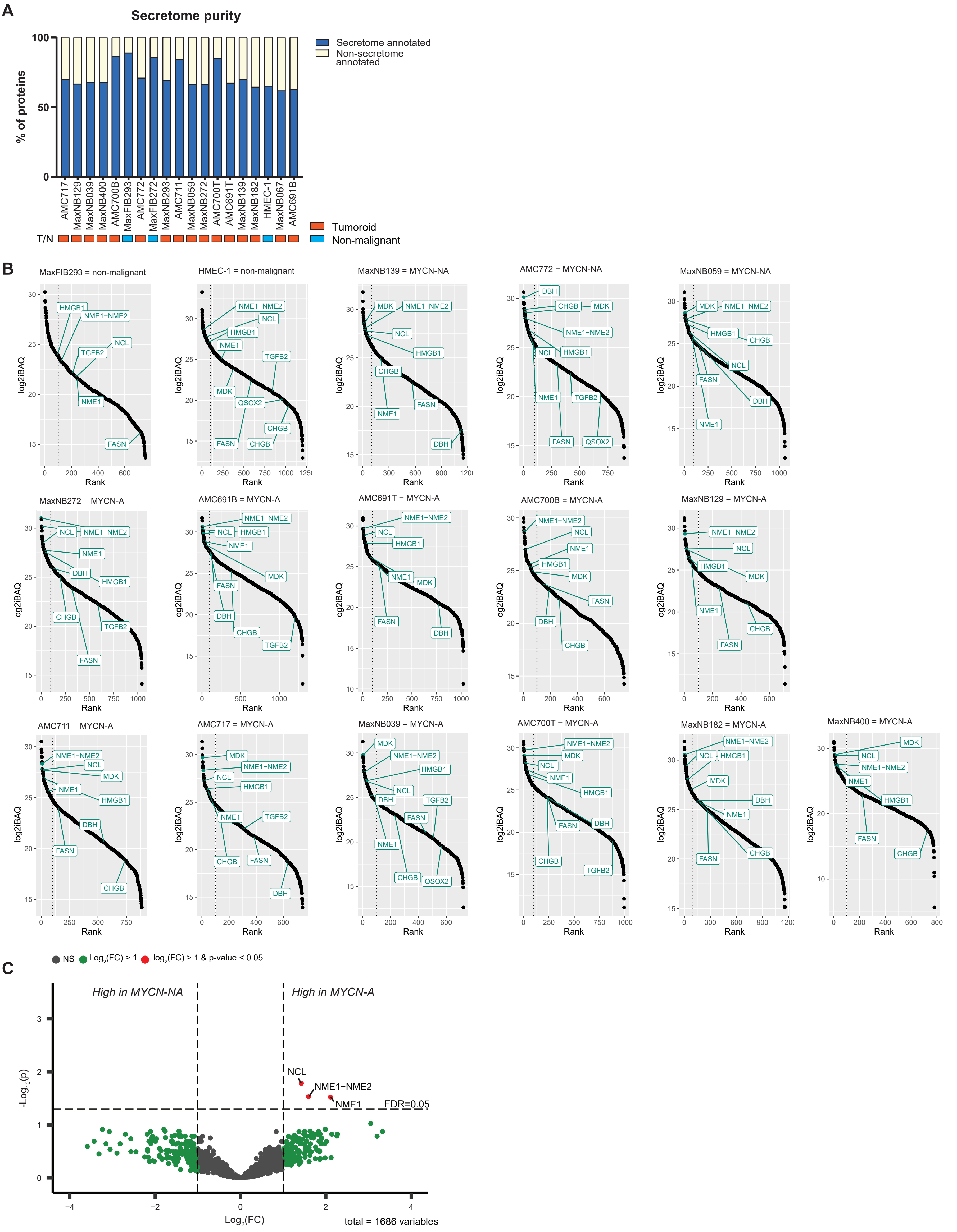

Supplementary figure 3

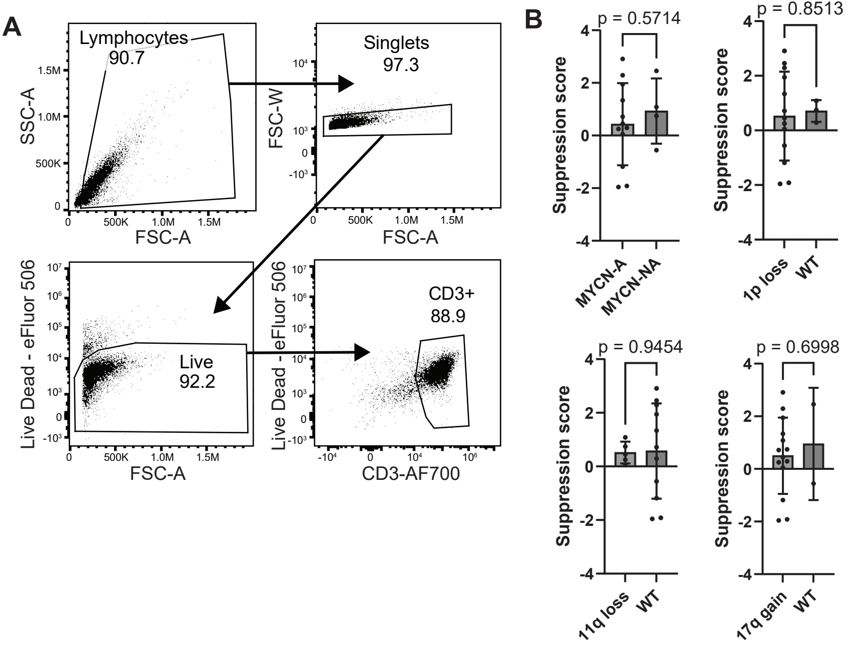

### Supplementary figure 4

**A**

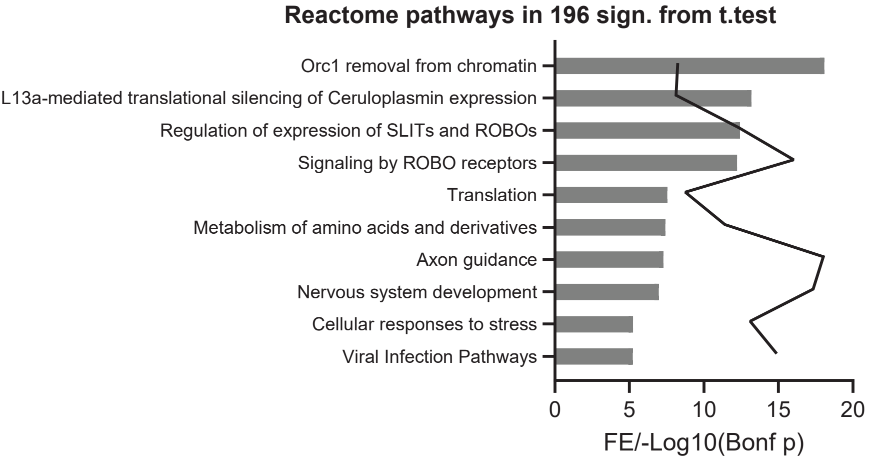

### Supplement Figure 5

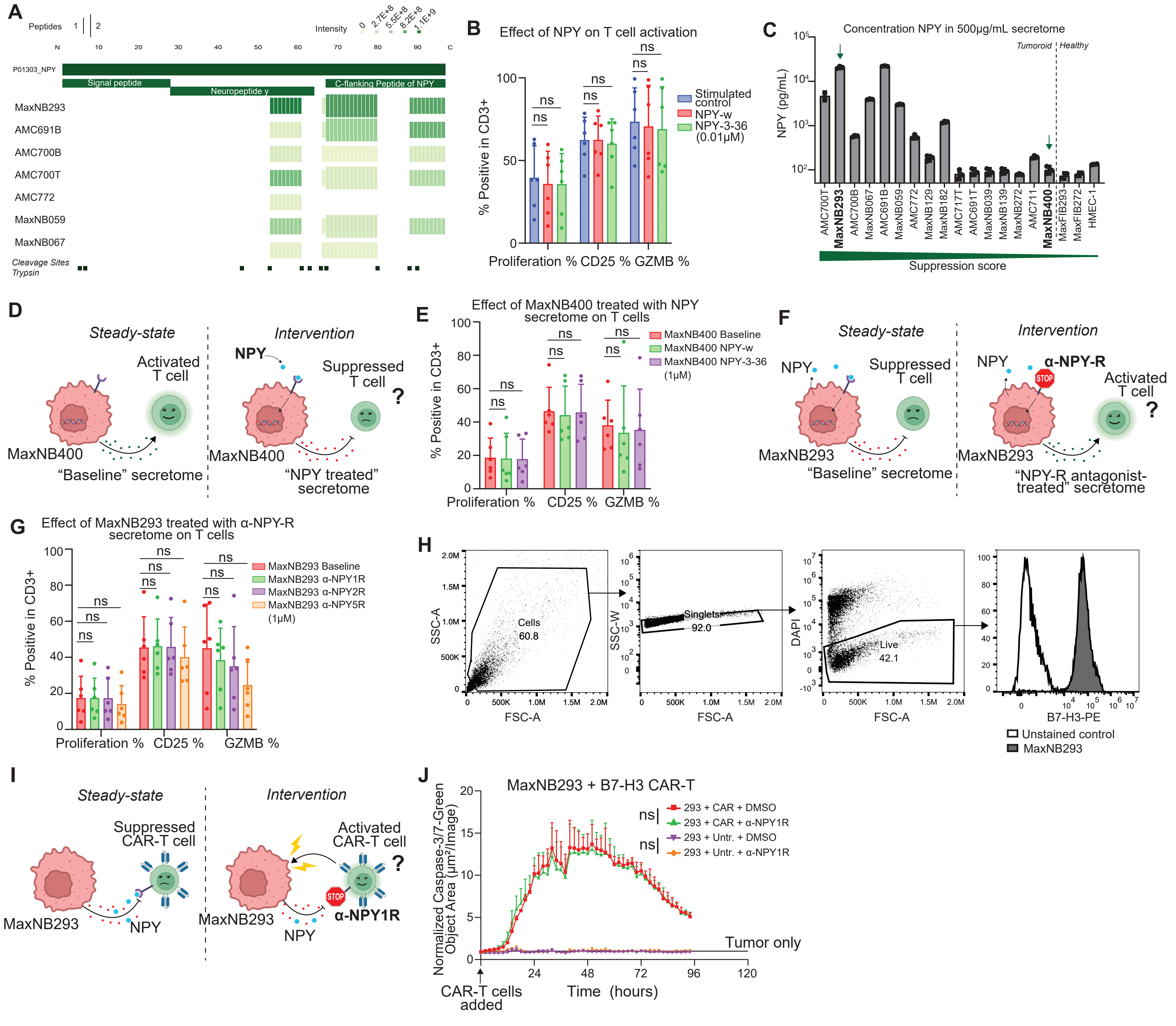

### Supplement Figure 6

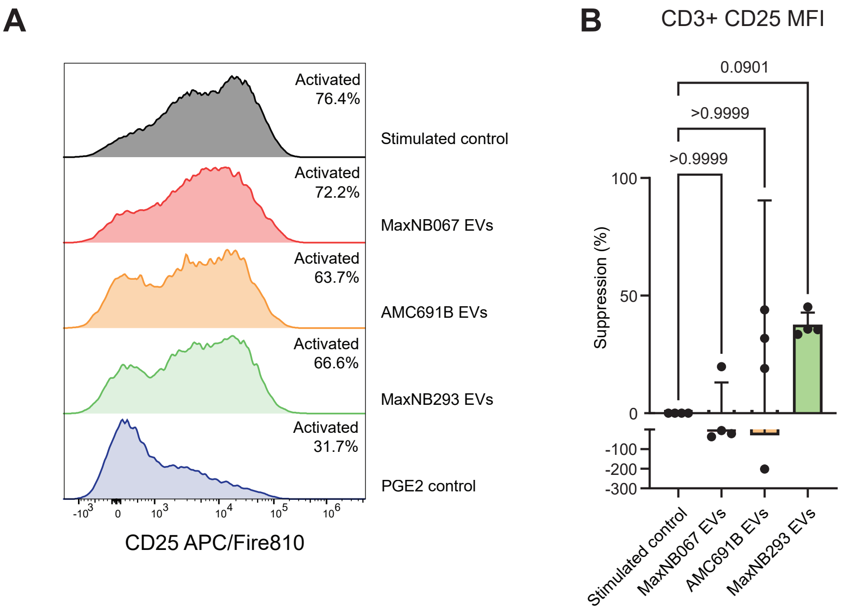

Supplementary Table 1

| Tumoroid name | Picture after conditioning | Cell count per T175 | Protein content (µg/mL) |
|---|---|---|---|
| MaxNB400 | 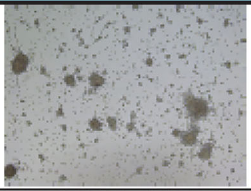 | 7.78E+06 | 4193 |
| MaxNB182 | 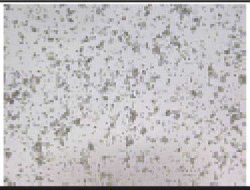 | 1.27E+07 | 3395 |
| MaxNB272 | 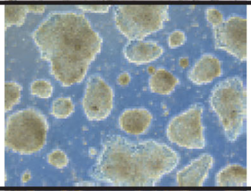 | 1.92E+07 | 7382 |
| MaxNB293 | 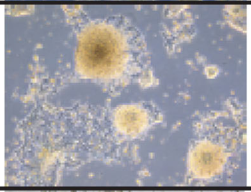 | 7.00E+06 | 6570 |
| AMC691B | 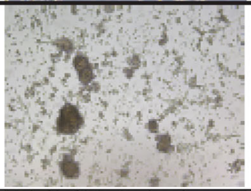 | 1.19E+07 | 6141 |
| AMC691T | 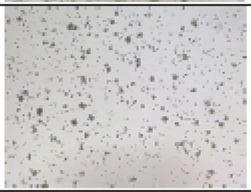 | 3.01E+07 | 3636 |
| AMC700B | 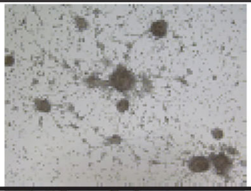 | 9.82E+05 | 1765 |
| AMC700T | 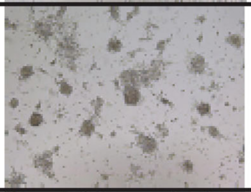 | 6.16E+06 | 2471 |
| AMC711 | 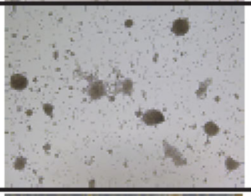 | 5.05E+06 | 3328 |
| AMC717 | 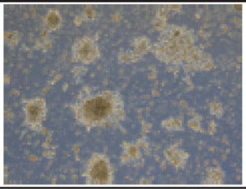 | 1.05E+07 | 4285 |
| AMC772 | 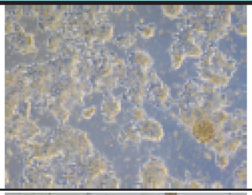 | 2.11E+07 | 11375 |
| MaxNB039 | 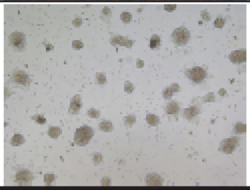 | 2.29E+07 | 6837 |
| MaxNB059 | 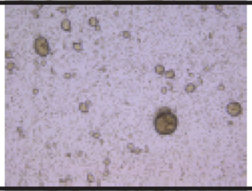 | 2.70E+06 | 5322 |
| MaxNB067 | 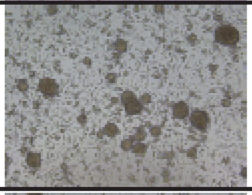 | 1.38E+07 | 5205 |
| MaxNB129 | 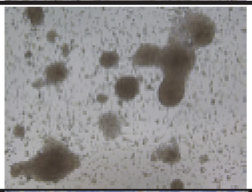 | 3.36E+06 | 2989 |
| MaxNB139 | 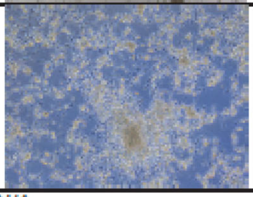 | 2.01E+07 | 6816 |
| 000DBE (Fibroblasts) | N/A | 2.35E+06 | 1760 |
| 000JEA (Fibroblasts) | N/A | 4.00E+05 | 1665 |
| HMEC-1 (Endothelial cell line) | 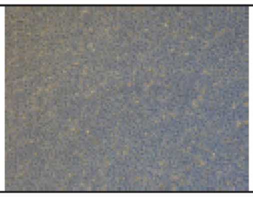 | 1.25E+07 | 483 |
